# Social and non-social autism symptom and trait domains are genetically dissociable

**DOI:** 10.1101/228254

**Authors:** Varun Warrier, Roberto Toro, Hyejung Won, Claire S Leblond, Freddy Cliquet, Richard Delorme, Ward De Witte, Janita Bralten, Bhismadev Chakrabarti, Anders D Børglum, Jakob Grove, Geert Poelmans, the 23andMe Research Team, David A. Hinds, Thomas Bourgeron, Simon Baron-Cohen

## Abstract

The core diagnostic criteria for autism comprise two symptom domains – social and communication difficulties, and unusually repetitive and restricted behaviour, interests and activities. There is some evidence to suggest that these two domains are dissociable, yet, this hypothesis has not been tested using molecular genetics. We test this using a GWAS of a non-social autistic trait, systemizing (N = 51,564), defined as the drive to analyse and build systems. We demonstrate that systemizing is heritable and genetically correlated with autism. In contrast, we do not identify significant genetic correlations between social autistic traits and systemizing. Supporting this, polygenic scores for systemizing are significantly positively associated with restricted and repetitive behaviour but not with social difficulties in autistic individuals. These findings strongly suggest that the two core domains of autism are genetically dissociable, and point at how to fractionate the genetics of autism.

## Introduction

The core diagnostic criteria of autism comprises two symptom domains: difficulties in social interactions and communication (the social domain) and unusually repetitive and restricted behaviour and stereotyped interests (the non-social domain)^1^. Multiple lines of evidence suggest that these two domains are dissociable^2,3^. First, factor and principal component analysis of autism and autistic traits have mostly identified two factors for autism – a social and a non-social factor^4–9^. Second, investigation of autistic traits in large cohorts have demonstrated a positive phenotypic correlation between different social traits and different non-social traits separately, but only a limited correlation between social and non-social traits^9–12^. Third, twin genetic correlations between social and non-social symptom domains in autism are low, though both social and non-social trait domains are highly heritable in neurotypical^13,14^ or autistic twins^15^. Fourth, difficulties in social and non-social domains can occur independently of each other^16,17^, which has been used to subgroup individuals on the spectrum based on the two domains^18^. This suggests that the genetic and phenotypic architecture of autism consists of at least two categories of broadly dissociable domains. This has implications for genetic, biological, and clinical studies of autism, since most studies have investigated autism as if it is a unitary condition^3^. The idea that social and non-social symptom domains are dissociable is unsurprising given their very different nature, and very different underlying cognitive processes one related to interpreting animate motion and mental states (theory of mind) and the other related to recognizing inanimate objects, events or patterns (systemizing), or their very different underlying neurology^3^. Nevertheless, the traditional view of autism is that it is a syndrome, meaning the diagnosis is only given when the social and non-social symptom domains cluster together.

However, to date, there has been limited molecular genetic evidence in support of this dissociability hypothesis, partly due to the limited large-scale research on the genetics of social and non-social domains. Most genetic research into the social and non-social domains has been primarily through linkage and genome-wide association studies (GWAS) in relatively small samples of autistic individuals and the general population (N < 5K)^19–25^. This has precluded a detailed molecular genetic investigation of the social and non-social domains associated with autism. Given currently available sample sizes with phenotypic information, investigating the genetics of the social and non-social domains in autistic individuals is difficult. However, several studies have demonstrated that the underlying liability for autism is normally distributed in the general population^26–29^. Factor analysis have failed to identify discontinuities between clinical autism and autistic traits in the general population^30^. Autistic traits are heritable^31–33^, are elevated in family members of autistic individuals compared to the general population^34,35^, and are transmitted intergenerationally^36,37^. Factor analysis of autistic traits measures have also identified two different factors in both the general population and autistic individuals – one linked to the social domain, and another linked to the non-social domain, mirroring the factor structure of clinical autism domains^6,9,30,38^. Studies have further demonstrated moderate to high shared genetics between the extremes of the liability distribution and the rest of the distribution^14,39–41^. One twin study investigated the bivariate genetic correlation between research and clinical autism diagnosis and autistic traits and identified high genetic correlations (0.7 < r_g_ < 0.89)^42^. Validating this, studies have identified modest shared genetics between autism and autistic traits^43–45^. Taken together, there is considerable evidence to suggest that autism represents the extreme end of the autistic traits continuum.

While a few studies have investigated the genetics of traits contributing to the social domains such as social and communications difficulties^19,44,45^, empathy^46^, and emotion recognition^47^, there have been limited studies investigating the genetics of the non-social domain^25,48^. Neither of these studies have replicably identified significant variants associated with the non-social domain, primarily because of the relatively modest sample sizes of the GWAS. An alternate approach is to investigate the genetics of non-social autistic traits in the typical population, maximizing the sample size. To better understand the genetics of a non-social autistic trait, we investigate the genetics of systemizing measured using a 75-item well validated self-report measure called the Systemizing Quotient-Revised (SQ-R) (**Methods**). Systemizing involves identifying *input-operation-output* relationships in order to analyse and build systems, and to understand the laws that govern specific systems^49^. The hyper-systemizing theory of autism proposes that autistic individuals, on average, have superior attention to detail, and a stronger drive to systemize^49^. This has been validated in several studies^50,51^ including a recent study in more 650,000 individuals including 36,000 autistic individuals^12^. Several lines of evidence suggest that autistic individuals have intact or superior systemizing. The idea was noted in the earliest papers describing autism by both Hans Asperger^52^ and Leo Kanner^53^. Further, autistic adults, on average, score higher on the SQ-R compared to individuals in the general population^10,51^, a pattern also observed in autistic children^54^. Several items in the SQ-R specifically measures circumscribed interests and insistence on sameness, two of the items mentioned in the DSM-5, and several of these items map onto items on the Autism Spectrum Quotient (AQ), a well validated measure of autistic traits^27^ (**Supplementary Note**). Because systems follow rules, they repeat, such that an operation on a given input produces the same output every time. A fascination with systems may thus manifest as unusually repetitive behaviour. And because systems depend on precise variables, a fascination with systems may also manifest as unusually narrow interests in autism.

The present study has two aims: 1. To investigate the polygenic architecture of a non-social trait linked to autism: *systemizing*, and 2. To investigate if social and non-social autistic traits measured in the general population are genetically dissociable.

## Results

We first conducted a GWAS of systemizing (N = 51,564) measured using the SQ-R. Following this, and using data from GWAS of social traits genetically correlated with autism (GWAS of self-reported empathy (N = 46,861)^46^, and GWAS of social relationship satisfaction^55^ measured using friendship (N_effective_ = 164,112) and family relationship (N_effective_ =158,116) satisfaction scales) we investigated if the social and non-social domains autistic traits are genetically dissociable in the general population. A flow-chart of the study design is provided in Figure 1.

**Figure 1.**
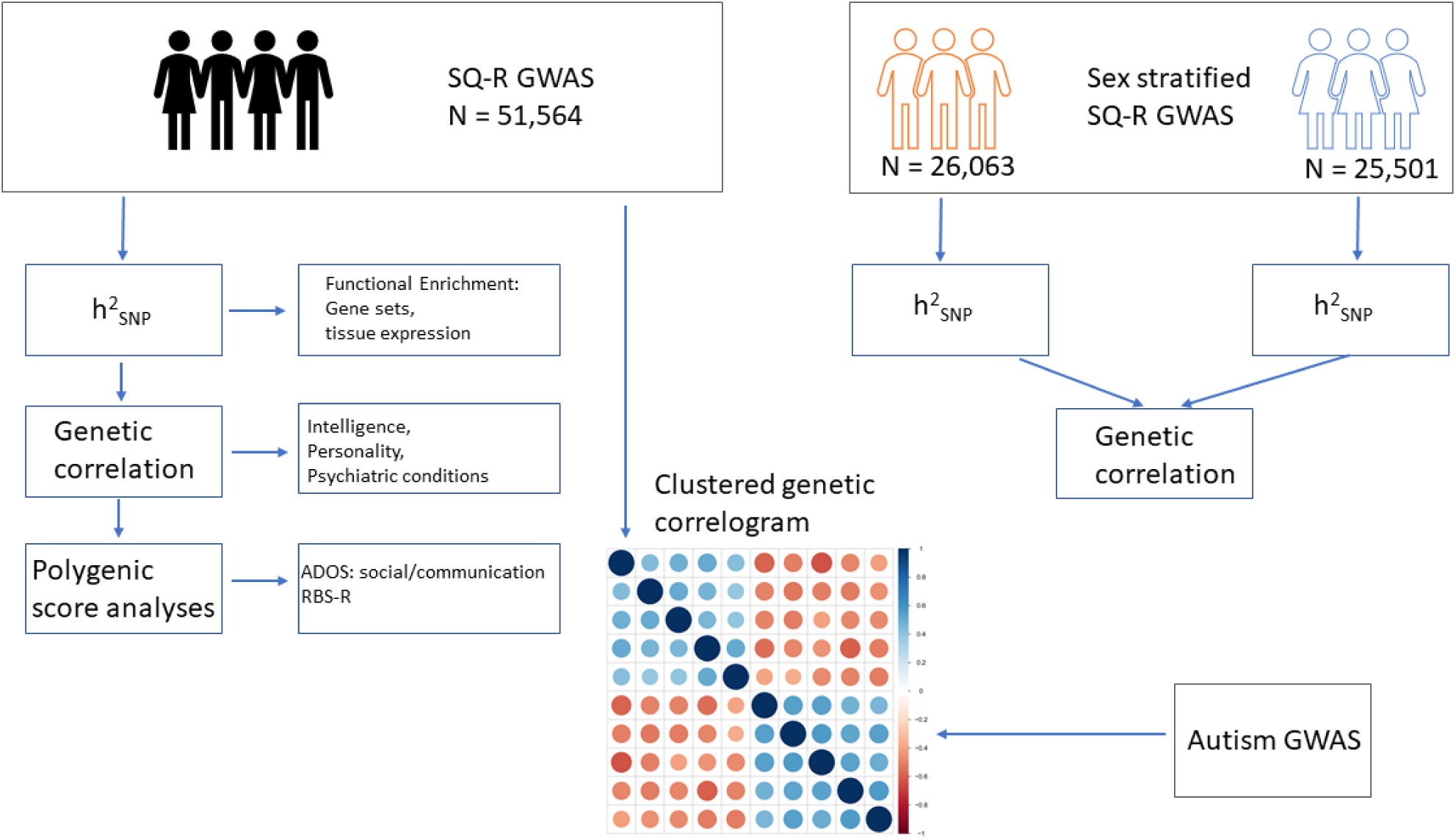
Schematic diagram of the study. We conducted a GWAS of the SQ-R (N = 51,564) and quantified SNP heritability (h^2^_SNP_), quantified genetic correlations with multiple phenotypes, and conducted polygenic score analyses. Additionally, we conducted sex-stratified GWAS of the SQ-R, and investigated h^2^_SNP_ within sex and genetic correlation between sex. Finally, we investigated the clustering of all phenotypes that are genetically correlated with autism, and if the social and the non-social phenotypes associated with autism are genetically correlated.

Systemizing was measured in the 23andMe sample (N = 51,564) using scores from the SQ-R^10^. Scores on SQ-R were normally distributed, with a mean of 71±21 out of 150. As hypothesized based on previous research^10,12,51^, males (76.5±20), on average, scored higher than females (65.4±20.6) (P < 0.001, Cohen’s d = 0.54, **Supplementary Figure 1**). Given the significant sex differences in scores, we conducted a non-stratified and sex-stratified GWAS for SQ-R. Genome-wide association analyses identified three significant loci (Figure 2, **Supplementary Table 1** and **Supplementary Figure 2**). Two of these were significant in the non-stratified GWAS: rs4146336 on chromosome 3 (P = 2.58×10^−8^) and rs1559586 on chromosome 18 (P = 4.78×10^−8^). The third significant locus was in the males-only GWAS (rs8005092 on chromosome 14, P = 3.74×10^−8^). rs8005092 and rs1559586 lie in regions of high genetic recombination. Linkage-Disequilibrium Score Regression (LDSR) intercept suggested that there was minimal inflation due to population stratification (Figure 2). Fine-mapping of the three regions identified 14 credible SNPs **(Methods)**. None of the SNPs overlapped with foetal brain eQTL. However, two of these SNPs mapped onto two genes - *LSAMP* and *PTMAP8*, both of chromosome 3 - using chromatin interaction data in the foetal brain. Of these, LSAMP is a neuronal adhesion molecule in the limbic system of the developing brain Additionally, gene-based analysis identified 4 significant genes *SDCCAG8*, *ZSWIM6*, *ZNF574* and *FUT8* **(Supplementary Table 2)**. Of these, mutations in *ZSWIM6* cause a neurodevelopmental disorder with, in some cases, co-morbid autism and unusually repetitive movements and behaviour^56^. As supporting analyses, we investigated the direction of effect for all independent SNPs with P < 1×10^−6^ in the non-stratified SQ-R GWAS in GWAS of autism^57^, educational attainment^58^, and cognitive aptitude^59^. Five out of six SNPs tested had concordant effect in the GWAS for educational attainment and GWAS for cognitive aptitude (P = 0.21, two-sided binomial sign test for each comparison). Similarly, four out of five SNPs tested had concordant effect direction in the GWAS for autism **(Supplementary Table 3a)** (P = 0.37, two-sided binomial sign test). For these three phenotypes, we additionally assessed effect direction concordance using binomial sign test at less stringent P-value thresholds in the SQ-R GWAS, after LD-based clumping (P < 1, 0.5, 0.1 and 1×10^−4^). Binomial sign test was statistically significant at three of the four P-value thresholds (P = 1, 0.5 and 0.1) for all three phenotypes but not statistically significant at P = 1E-4, presumably due to the low statistical power (**Supplementary Table 3b**). Additionally, we tested effect direction concordance (P < 1×10^−6^) in a GWAS (N = 1,981) of ‘insistence on sameness’, a phenotype that’s similar to systemizing (**Methods**). Four out of five SNPs had a concordant effect direction including the two SNPs with P < 5×10^−8^ in the non-stratified SQ-R GWAS (P = 0.37, two-sided binomial sign test).

**Figure 2:**
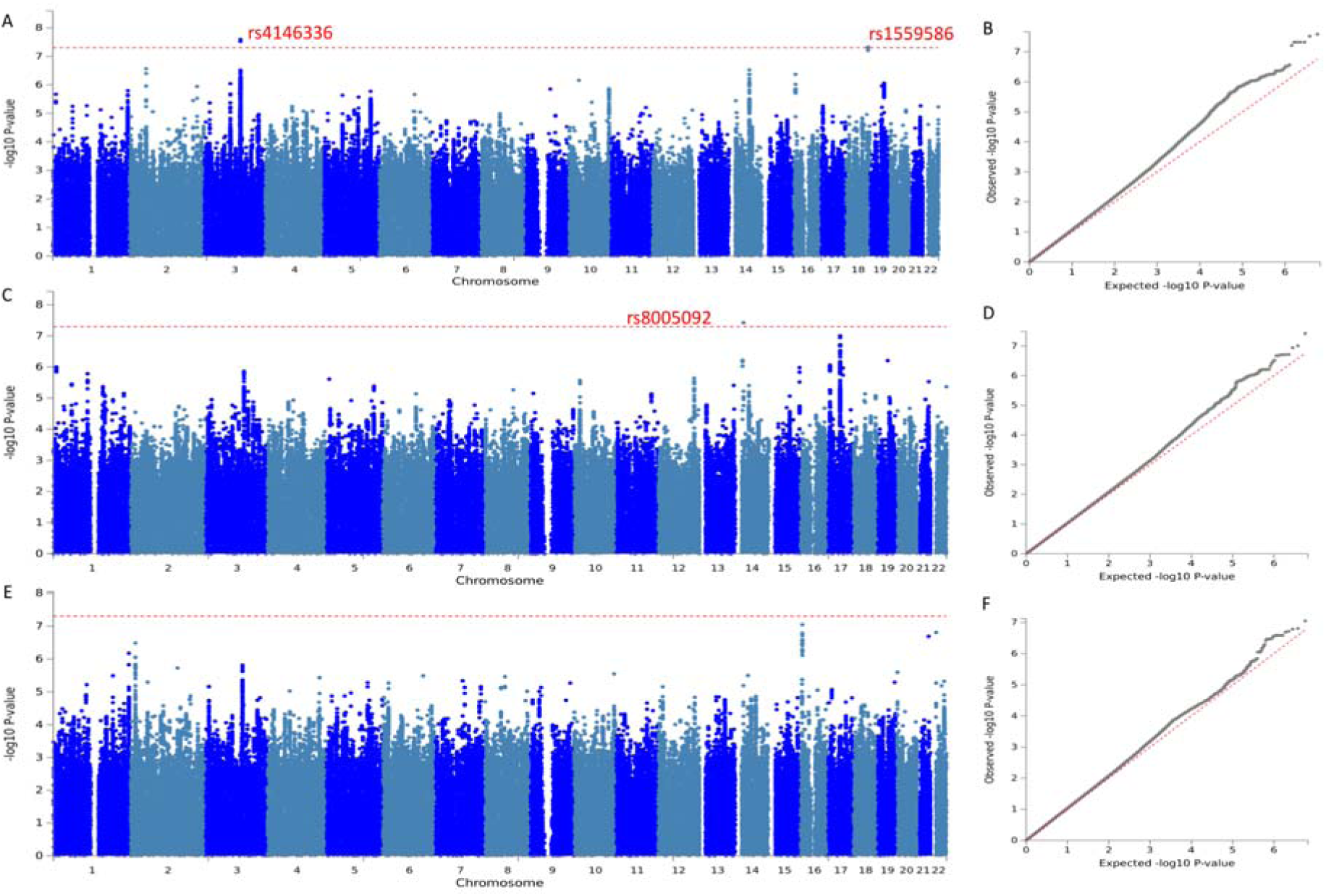
Manhattan and QQ-plots for the three GWAS. Manhattan plot for the three SQ-R GWAS: non-stratified (A), males-only (C), females-only (E). Significant SNPs are highlighted in red. QQ-plots for the three SQ-R GWAS: non-stratified (B), males-only (D), females-only (F). SQ-R non-stratified (N = 51,564): λ_GC_ = 1.10, LDSR intercept = 0.99, SQ-R males only (N = 26,063): λ_GC_ = 1.06, LDSR intercept = 0.99, SQ-R females only (N =25,501): λ_GC_= 1.05, LDSR intercept = 1.01.

Additive SNP-based heritability (h^2^_SNP_) calculated using LDSR was 0.12±0.012 for the SQ-R (P = 1.2×10^−20^). Despite small but significant sex-differences in the SQ-R scores, there was no significant difference in h^2^_SNP_ between males and females (P = 0.34) **(Supplementary Figure 3 and Supplementary Table 4)**, which was strengthened by the high genetic correlation between males and females (1±0.17; P = 3.91×10^−10^), suggesting a similar polygenic architecture between sexes. The per-SNP effect for the most significant SNPs was small, suggesting a highly polygenic architecture (R^2^ = 0.001 – 0.0002%, after correcting for winner’s curse, **Supplementary Table 5**).

Partitioned heritability for functional categories identified significant enrichment for evolutionary conserved regions, transcription start sites, foetal DNase hyper-sensitivity sites, and H3 lysine 27 acetylation (H3K27ac), suggesting a prominent role for regulatory and conserved genomic regions in systemizing **(Supplementary Table 6)**. Partitioning heritability based on tissue specific active chromatin marks identified a significant enrichment for brain specific chromatin signatures highlighting the role of the brain for the SQ-R. Notably, this enrichment was significant in both adult and foetal brain specific active chromatin marks **(Supplementary Table 7 and Supplementary Figure 4)**. Enrichment for genes expressed in the brain was high but failed to reach statistical significance after correcting for the multiple tests conducted (**Supplementary Figure 5 and Supplementary Table 8**).

We identified a significant positive genetic correlation between the SQ-R and autism as well as measures of intelligence (cognitive aptitude and educational attainment) **(Supplementary Table 9 and** Figure 3a). Of all the psychiatric conditions tested (**Methods**), SQ-R was only significantly genetically correlated with autism (r_g_ = 0.26±0.06; P = 3.35×10^−5^), demonstrating the relative specificity of the SQ-R to autism. Notably, the effect size of the genetic correlation between autism and the SQ-R is similar to the genetic correlation between autism and self-reported empathy (measured using the Empathy Quotient (EQ^60^): r_g_ = −0.27 ± 0.07) and scores on the Social and Communication Disorders Checklist (SCDC^61^): r_g_ = 0.27 ± 0.13). Controlling for the genetic effects of educational attainment on the SQ-R GWAS using genome-wide inferred statistics (GWIS) (**Methods**) attenuated the genetic correlation with autism only modestly, suggesting that the SQ-R scores are genetically correlated with autism independently of the genetic effects of education (Figure 3b **and Supplementary Table 10)**. We validated this using genomic structural equation modeling (GSEM) (**Methods**) using both educational attainment and cognitive aptitude (Figure 3c). Further, the SQ-R was not genetically correlated with any of the social measures related to autism – friendship and family relationship satisfaction, scores on a self-report measure of empathy - the Empathy Quotient (EQ), and the scores on the Social and Communication Disorders Checklist (SCDC), which is a measure of social and communication difficulties (see **Supplementary Note** for how these traits map onto social domains in autism). Estimates of genetic correlations between SQ-R scores and the various social traits are also small, suggesting that there is limited shared genetics between social autism traits and the SQ-R.

**Figure 3:**
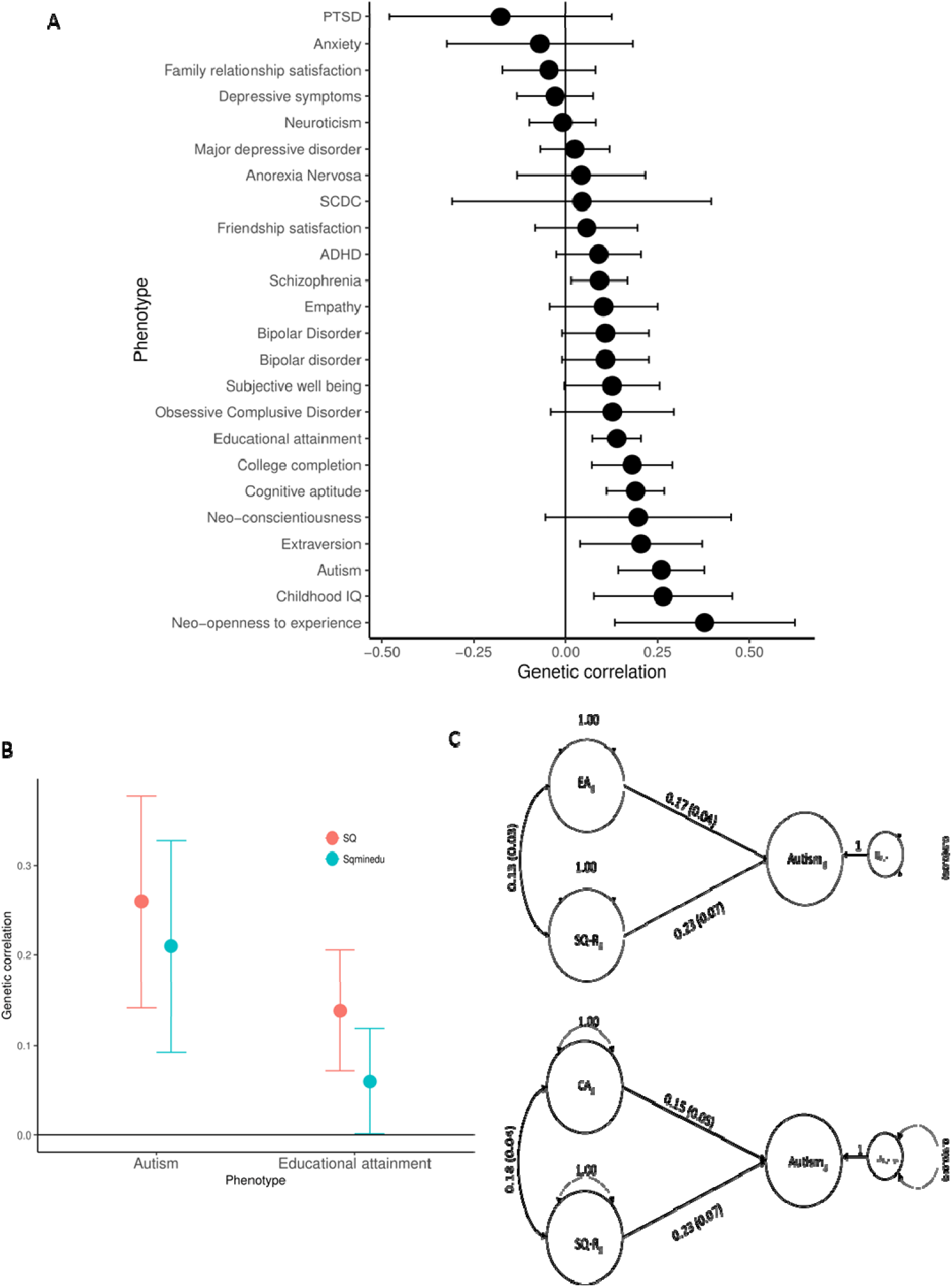
Genetic correlation between the SQ-R and other phenotypes, and GWIS and GSEM estimates between SQ, educational attainment and cognitive aptitude. 3A: Genetic correlations between the SQ-R and multiple other phenotypes provided. The bars represent 95% confidence intervals. Sample sizes and PMID are provided in Supplementary Table 9. The following genetic correlations were significant after Bonferroni correction: Autism (r_g_ = 0.26±0.06; P = 3.35×10^−5^, N = 46,350), Years of Schooling 2016 (r_g_ = 0.13±0.03; P = 4.73×10^−5^, N = 293,723), College completion (r_g_ = 0.18±0.05; P = 1.30×10^−3^, N = 95427), and Cognitive aptitude (r_g_ = 0.19±0.04; P = 2.35×10^−5^, N = 78,308). 3B: Results of the GWIS analysis. Red lines represent genetic correlation with the SQ-R, blue lines represent genetic correlations with the SQ-R independent of the genetic effects of educational attainment. The bars represent 95% confidence intervals. 3C: Path diagrams providing the results of the standardized SEM models to investigate if the SQ-R is genetically correlated with autism independent of the genetic effects of cognitive aptitude (CA_g_) and educational attainment (EA_g_).GWIS is Genome-wide inferred statistics; GSEM is Genomic structural equation modeling.

To understand the genetic relationship between the SQ-R and autism in a broader context, we evaluated the genetic correlations between multiple phenotypes with evidence of significant genetic correlation with autism (15 phenotypes in total, see **Methods** for a list of phenotypes included). Clustering highlighted three broad clusters: a social cluster, a psychiatric cluster, and an intelligence cluster (Figure 4a **and Supplementary Tables 11 and 12)**. The SQ-R clusters closely with measures of intelligence, but while educational attainment and cognitive aptitude are significantly correlated with multiple social traits and psychiatric conditions, the SQ-R is only genetically correlated with autism.

**Figure 4:**
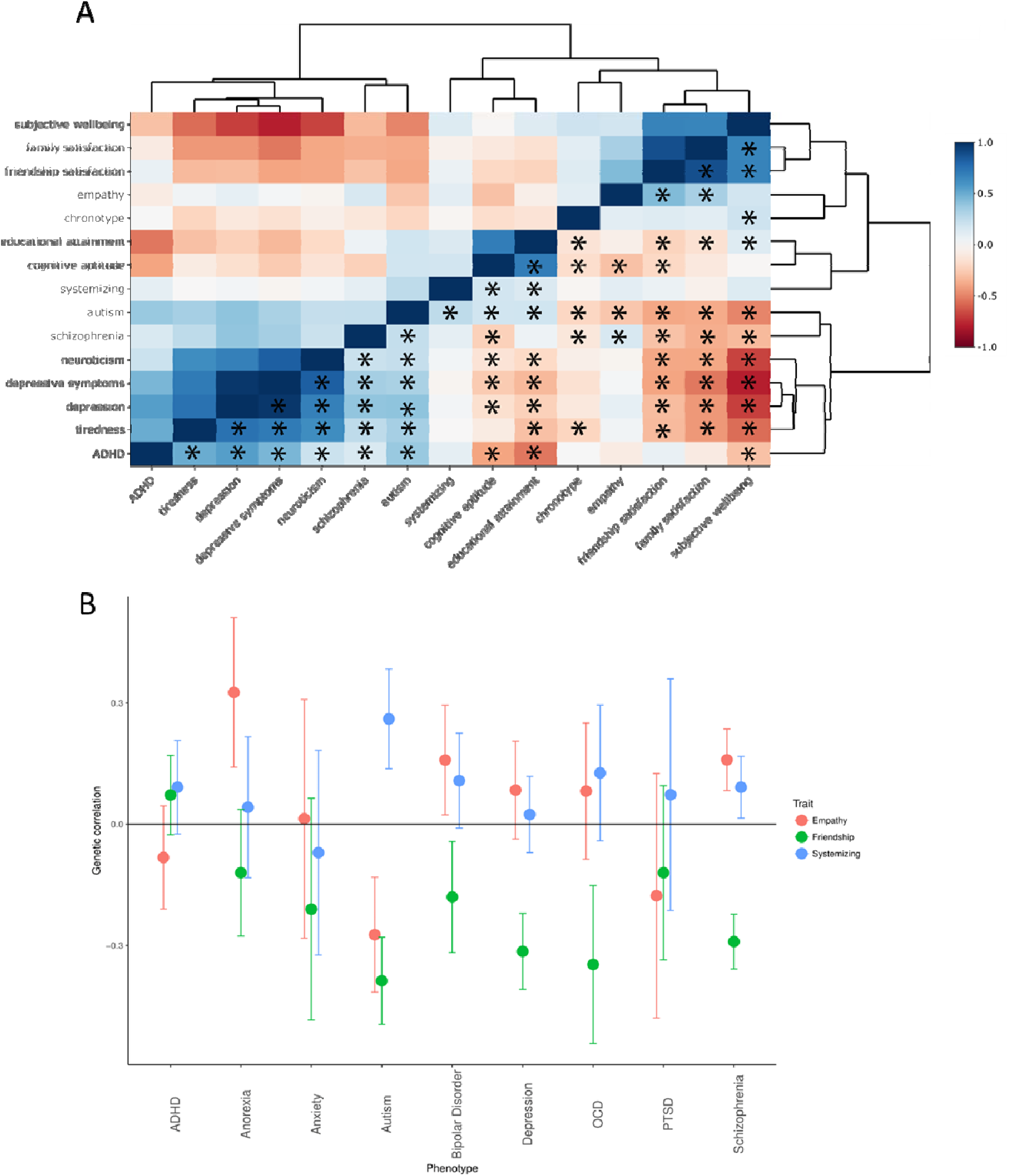
Genetic correlogram of autism and related traits, and genetic correlations between social and non-social traits and multiple psychiatric conditions. 4A: Correlogram of genetic correlations between all phenotypes that are genetically correlated with autism. Please note the upper and lower triangle are identical. Asterisk (provided only in the lower triangle) represent significant correlations after Bonferroni correction. Genetic correlations have been clustered using hierarchical clustering. Colour as provides the magnitude of genetic correlation. 4B: Genetic correlation between empathy, friendship satisfaction, and systemizing with nine psychiatric conditions. Only autism was significantly genetically correlated with all three phenotypes.

Given that the two major domains of autism as identified by the DSM-5 are persistent difficulties in social interaction and communication and unusually restrictive, stereotyped, and repetitive interests^1^, we hypothesized that the combination of significant negative genetic correlation with social traits (friendship satisfaction and empathy) and significant positive genetic correlation with SQ-R is uniquely associated with autism (**Methods**). Indeed, across the nine psychiatric conditions for which we had summary GWAS statistics, this combination was uniquely seen for autism (Figure 4b, **Supplementary Table 13)**.

Given that our current analysis focussed on the general population, we sought to investigate if polygenic scores from the SQ-R were associated with social and non-social autism domains in 2,221 autistic individuals from the Simons Simplex Collection (**Methods**). We hypothesized that SQ-R may be significantly associated with the non-social domain in autism, but not associated with the social domain in autism. Polygenic scores for SQ-R were significantly associated with scores on the Repetitive Behaviour Scale-Revised (Beta = 0.052±0.02, P = 0.013), but not on the social and communication subscale of ADOS-G (Beta = −0.00099±0.018, P = 0.95) after adjusting for multiple test (Bonferroni alpha = 0.025). We validated this in 426 additional individuals of which 401 had a diagnosis of autism with RBS-R scores from the EU-AIMS LEAP, AGRE, and Paris cohorts. Here, we identified a concordant effect direction for polygenic score of the SQ-R (Beta = 0.02±0.05, P = 0.65), though the results were not significant potentially due to the small sample size. Inverse-variance meta-analysis of the discovery and the validation cohorts marginally improved the significance of the association (Beta = 0.047±0.018, P = 0.010), and the results remained statistically significant (Bonferroni alpha = 0.025). In a separate sample of 475 autistic individuals from the AGRE cohort, polygenic scores for the SQ-R were not associated with the social and communication subscale of ADOS-G (Beta = −0.046±0.04, P = 0.24). Meta-analysis of the two cohorts did not produce a statistically significant result (Beta = - 0.008±0.016, P = 0.60) (see **Power calculations in the Supplementary Note)**. We note that the lack of association between the polygenic scores for the SQ-R and the ADOS-G social and communication subscale is not indicative of absence of shared genetics, but rather indicative of lower shared genetics than between the RBS-R and the SQ-R.

Finally, to further validate the results in autistic individuals, we conducted bivariate genetic correlations on scores on the RBS-R and the ADOS-G social and communication subscale in 2,989 individuals from the SSC, AGRE, EU-AIMS LEAP and Paris cohorts (2964 autistic individuals). Both the RBS-R (h^2^_SNP_ = 0.11±0.11, P = 0.15) and the ADOS-G social and communication subscale (h^2^_SNP_ = 0.26±0.10, P = 0.004) had modest h^2^_SNP_, though only the latter was statistically significant. We identified a small genetic correlation (r_g_ = 0.15±0.46, P = 0.74), which was not statistically different from 0. Given the small sample size, the genetic correlation is unlikely to be statistically significant. However, the effect was small and statistically less than 1 (P = 0.034, One-tailed T test).

## Discussion

We present the largest GWAS of a non-social trait related to autism in the general population – systemizing, measured using the SQ-R. We demonstrate that systemizing is heritable and genetically correlated with autism. Associated loci are enriched in genomic regions containing brain chromatin signatures and we identify three genome-wide significant loci, but these must be replicated in an independent cohort. Despite the modest sample size, our GWAS is well-powered to investigate genetic correlations between various phenotypes including social traits related to autism, as the Z-score of the h^2^_SNP_ is above the recommended threshold of four^62^. We identify high sign concordance of the top SNPs in genetically correlated traits, enrichment for active chromatin marks in foetal and adult brain, and significant polygenic score association with the RBS-R. Polygenic score analysis suggests that the shared genetics between systemizing and the non-social domain in autism is considerably higher than the shared genetics between systemizing and the social domain. In addition, using a smaller sample of autistic individuals, we provide preliminary evidence that the social and non-social domains in autistic individuals have low shared genetics. Our results highlight the need to collect deeper clinical and cognitive information in autistic individuals to better understand the phenotypic heterogeneity in autism.

Most studies model autism, and the underlying liability measured as autistic traits, as a single domain. This has likely arisen because of the difficulties in recruiting and phenotyping sufficient numbers of autistic people. Our study suggests that, both in the general population and in autistic individuals, social and non-social autistic traits and symptom domains are genetically dissociable. This may to some extent explain why, compared to GWAS of other psychiatric conditions of roughly similar sample sizes^57,63–65^, GWAS of autism to date have identified fewer loci. One possible explanation is statistical signal-attenuation because of the underlying heterogeneity. However, this does not necessarily suggest that systemizing, or the other individual trait domains are less complex. For instance, we observe similar h^2^_SNP_ for SQ-R, self-reported empathy ^46^, and the largest and most recent GWAS of autism^57^

It is important to investigate if these domains are dissociable in a larger cohort of autistic individuals and identify potential convergence of the two domains in gene expression networks in the developing brain. Our results confirm the need to rethink our understanding of autism as existing along a single dimension^3,66^. We hypothesize that the dissociation of the two domains will extend to these other research modalities in studies of autism and autistic traits. It is important to note that, while our results demonstrate two broadly dissociable autistic trait domains in the general population and in autistic individuals, more research is needed to identify other potentially dissociable domains and to investigate if this dissociability is driven by different designs of phenotypic instruments. For example, our research does not make a distinction between communication vs. social interaction abilities, or between sensory difficulties vs. repetitive behaviours, and future molecular genetic studies may identify varying levels of overlap between these domains. The same principle applies to other research modalities (neuroimaging, cognitive studies, hormonal assays, etc.,) investigating the biology of autism and autistic traits. These different symptom domains of autism may contribute to different co-morbidities. Our results identify shared genetics between the social autistic traits and psychiatric conditions such as schizophrenia and depression, but limited shared genetics between the SQ-R and these conditions. This needs to be evaluated epidemiologically.

## Methods

### Participants

The current study included participants from 23andMe (Primary GWAS - SQ-R), from ALSPAC (GWAS of scores on the Social and Communication Disorders Checklist (SCDC)) and autistic individuals from the Simons Simplex Collection (SSC), the Autism Genetic Resource Exchange (AGRE), and the EU-AIMS LEAP and PARIS cohorts.

### 23andMe

Research participants in the GWAS of the SQ-R were from 23andMe and are described in detail elsewhere^67,68^. All participants provided informed consent and answered surveys online according to a human subjects’ research protocol, which was reviewed and approved by Ethical & Independent Review Services, an external AAHRPP-accredited private institutional review board (http://www.eandireview.com). All participants completed the online version of the SQ-R on the 23andMe participant portal. Only participants who were primarily of European ancestry (97% European Ancestry) were selected for the analysis using existing methods^69^. Unrelated individuals were selected using a segmental identity-by-descent algorithm^70^. A total of 51,564 participants completed the SQ-R (males = 26,063, and females = 25,501).

### ALSPAC

ALSPAC is a longitudinal cohort which recruited pregnant mothers in the Avon region of the UK. The ALSPAC cohort comprises 14,541 initial pregnancies from women in Avon resulting in a total of 13,988 children who were alive at 1 year of age. Children were enrolled in additional phases, described in greater detail elsewhere^71^. This study received ethical approval from the ALSPAC Law-and-Ethics Committee, and the Cambridge Human Biology Research Ethics Committee. Written informed consent was obtained from parent or a responsible legal guardian for the child to participate. Assent was obtained from the child participants where possible. We conducted a GWAS of scores on the SCDC in 5,421individuals from ALSPAC.

### Other cohorts

We included data from four cohorts to conduct polygenic score and bivariate genetic correlation analysis. The SSC (n = 2,221 unrelated autistic individuals) consists of simplex autistic families, and are described elsewhere^72^. The AGRE cohort (n = 482 unrelated autistic individuals) consists of multiplex autism families, details of which are provided elsewhere^73^. Across all cohorts, all participants were of European ancestry as identified using multi-dimensional scaling. Additionally, we included 401 individuals (including 25 neurotypical individuals) from the EU-AIMS LEAP^74^ and Paris^75^ cohorts. Across all cohorts, we included only unrelated individuals, who were predominantly of European Ancestry as defined by genetic principal components (5 SD deviations above or below the mean European PC1).

Additionally, we also included data from 1,981 unrelated individuals (1000 males, 1981 females) from the Nijmegen Biomedical Study (NBS) to provide support for the independent SNPs with P < 1×10-6 in the non-stratified GWAS. Participants were asked the question: “It upsets me if my daily routine is disturbed”, which is related to a non-social domain of autism, and is similar to an item in the Autism Spectrum Quotient. Further information including genotyping and quality control is provided elsewhere^43^. Genetic association for the top SNPs were conducted using age, sex, and the first five genetic principal components as covariates using linear regression.

### Phenotypes

The primary phenotype for this study is the SQ-R, which was used to conduct a GWAS in participants from 23andMe. The SQ-R is self-report measure of systemizing drive, or interest in rule-based patterns^10^. The SQ-R taps a variety of domains of systemizing, such as interest in mechanical (e.g., car engines), abstract (e.g., mathematics), natural (e.g., the weather), motor (e.g., knitting), and collectible (e.g., stamp collecting) systems. There are 75 items on the SQ-R, with a maximum score of 150 and a minimum score of 0. Scores on the test are normally distributed^10^. The SQ-R has good cross-cultural stability and good psychometric properties with Cronbach’s alpha ranging from 0.79 to 0.94 in different studies^76^. Test-retest reliability available in a Dutch sample indicated high reliability of 0.79 (Pearson correlation)^76^. This was supported by another study in 4,058 individuals which identified high internal cohesion^77^. Exploratory followed by confirmatory factor analysis using Rasch modelling suggests that the SQ-R is unidimensional^77^. A sex difference has been observed in multiple studies with males, on average, scoring significantly higher than females^10,51^. Criterion validity shows that the SQ-R has a modest but significant correlation with the Mental Rotation Test (r =.25, P =.013), as well as its subscales^78^. Autistic individuals, on average, score higher on the SQ-R in multiple different studies^10,51,79^. Further, the SQ-R also predicts autistic traits, with a combination of the SQ-R and the Empathy Quotient predicting as much as 75% of the variance on the Autism Spectrum Quotient, a measure of autistic traits^10^. The SQ-R has been validated using a short form in a very large population of 600,000 controls and 36,000 autistic individuals (Greenberg et al, 2018).

In addition, we used the following secondary phenotypes: SCDC in ALSPAC, ADOS-G social and communication scores and the RBS-R in the other cohorts. We also used a single question which is a measure of ‘insistence on sameness’ in the NBS cohort.

The SCDC is a questionnaire that measures difficulties in verbal and nonverbal communication, and social interaction including reciprocal social interaction^61^. The questionnaire consists of 12 questions, with scores ranging from 0 – 24, with higher scores reflecting difficulties in social interaction and communication. The SCDC has good internal consistency (0.93) and good test-retest reliability (0.81)^61^. The SCDC has reasonable specificity and sensitivity in distinguishing clinically diagnosed autism from control individuals^80^. Previous research has demonstrated that the SCDC is genetically correlated with autism^44,45,57^. We conducted a GWAS of SCDC to investigate if it is genetically correlated with SQ-R in this study. We used mother-reported SCDC scores on children aged 8. While SCDC has been measured at different ages in the ALSPAC cohort, we chose SCDC scores at age 8 as these had the largest sample size and have high h^2^_SNP_ ^19^ (h^2^ = 0.24 ± 0.07).

We chose two measures of social and non-social traits. For the social trait, we used the social and communication domain scores from the ADOS-G, a widely used instrument for diagnosing and assessing autism in four cohorts (SSC, AGRE, EU-AIMS LEAP and Paris). Participants completed one of the following ADOS-G modules^81^: 1 (used for children with little or no phrase speech), 2 (for children with non-fluent speech), 3 (verbally fluent children), and 4 (verbally fluent adolescents and adults). For this study, we used the raw totals of the scores from the social domain and the communication domain, combined. Scores for all 4 modules range from 0 – 24. The ADOS-G has high overall internal consistency, and high test-retest reliability for the social and communication subscales^81^. The choice for combining the social and communication domain scores were informed by factor analysis which suggested that the two domains contribute to one underlying factor^82^.

In contrast to the Social and Communication domain, the restricted and repetitive behaviour domain of the ADOS-G has poor test retest reliability (r < 0.6) and a smaller range of scores (0 – 8) as it captures fewer repetitive and restrictive behaviour^81^. Hence, for this study, we used sores on the RBS-R^83^. The RBS-R is a measure developed to specifically measure restricted and repetitive behaviours in autistic individuals and captures stereotyped, self-injurious, sameness, compulsive, ritualistic, and restricted behaviour^84^, and has high inter-rater reliability and internal consistency^84^. The RBS-R comprises 43 questions with scores ranging from 0 – 3 for each item based on a Likert scale.

‘Insistence on sameness’ in the NBS cohort was measured using a single item: “It upsets me if my daily routine is disturbed”. This is related to a non-social domain of autism, and is again similar to an item in the Autism Spectrum Quotient. Participants were asked to indicate on a 4-point Likert scale “definitely agree”, “slightly agree”, “slightly disagree”, “definitely disagree”.

### Genotyping, imputation, and quality control and genetic association in the 23andMe cohort

Details of genotyping, imputation and quality control in the 23andMe cohort are provided elsewhere^47^. Briefly, unrelated participants were included if they had a call rate of greater than 98.5%, and were of primarily European ancestry (97% European ancestry). A total of 1,030,430 SNPs (including InDels) were genotyped. SNPs were excluded if: they failed the Hardy-Weinberg Equilibrium Test at P < 10^−20^; had a genotype rate of less than 90%; they failed the parent-offspring transmission test using trio data in the larger 23andMe research participant database; or if allele frequencies were significantly different from the European 1000 Genomes reference data (chi-square test, P < 10^−20^). Phasing was conducted using Beagle (version 3.3.1)^85^ in batches of 8000-9000 individuals. This was followed by imputation against all-ethnicity 1000 Genomes haplotypes (excluding monomorphic and singleton sites) using Minimac2^86^. Genetic association analyses were restricted to SNPs with a minor allele frequency > 1%. After quality control, 9,955,952 SNPs (imputed and genotyped) were included in the GWAS.

Our primary analysis was an additive model of genetic effects and was conducted using a linear regression with age, sex, and the first five ancestry principal components included as covariates. In addition, given the modest sex difference, we also conducted sex-stratified analyses. SNPs were considered significant at a genome-wide threshold of P <5×10^8^. Leading SNPs were identified after LD-pruning using Plink (r^2^ > 0.8). Winner’s curse correction was conducted using an FDR based shrinking^87^.

We calculate variance explained by first standardizing the regression estimates and then squaring the estimates. This is equivalent to:

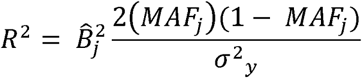

Where R^2^ is the proportion of variance explained for SNP j.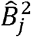 is the non-standardized regression coefficient, MAF is the minor allele frequency for SNP j, and *σ*^2^_*y*_ is the variance of SQ. Further details of this formula are provided in the **Supplementary Note**.

### Genotyping, imputation, and quality control and genetic association in the ALSPAC

The SCDC ^61^ scores were calculated from children of the 90s (ALSPAC cohort)^71^, in children aged 8. In total, SCDC scores were available on N = 7,825 children. From this, we removed individuals for whom complete SCDC scores were not available. After excluding related individuals and individuals with no genetic data, data was available on a total of N = 5,421 unrelated individuals.

Participants were genotyped using the Illumina® HumanHap550 quad chip by Sample Logistics and Genotyping Facilities at the Wellcome Sanger Institute and LabCorp (Laboratory Corportation of America) using support from 23andMe. Individuals were excluded based on gender mismatches, high missingness (> 3%), and disproportionate heterozygosity. We restricted subsequent analyses to individuals of European descent (CEU), which were identified by multidimensional scaling analysis and compared with Hapmap II (release 22). Individuals were also removed if cryptic relatedness, assessed using identity by descent, was greater than 0.1. Genotyped SNPs were filtered out if they had more than 5% missingness, violated Hardy-Weinberg equilibrium (P < 1×10^−6^), and had a minor-allele frequency less than 1%, resulting in a total of 526,688 genotyped SNPs. Haplotypes were estimated using data from mothers and children using ShapeIT (v2.r644)^88^. Imputation was performed using Impute2 V2.2.2^89^ against the 1000 genomes reference panel (Phase 1, Version 3). Imputed SNPs were excluded from all further analyses if they had a minor allele frequency < 1% and info < 0.8. After quality control, there were 8,282,911 genotyped and imputed SNPs that were included in subsequent analyses.

Dosage data from BGEN files were converted using hard-calls, with calls with uncertainty > 0.1 treated as missing data. Post-imputation, we excluded SNPs that deviated from Hardy-Weinberg Equilibrium (P < 1×10^−6^), with minor allele frequency < 0.01 and missing call rates > 2%. We further excluded individuals with genotype missing rates > 5%. The SCDC score was not normally distributed so we log-transformed the scores and ran regression analyses using the first two ancestry principal components and sex as the covariates using Plink 2.0^90^.

The log-transformed SCDC scores (henceforth, SCDC scores) had a modest but significant h^2^_SNP_ as quantified using LDSR (h^2^_SNP_ = 0.12 ±0.05). LDSR intercept (0.99) suggested that there was no inflation in GWAS estimates due to population stratification. The λ_GC_ was 1.013. We replicated the previously identified genetic correlation with autism^57^ (constrained intercept) using our SCDC GWAS (r_g_ = 0.45±0.18, P = 0.01). In addition, we also identified a negative genetic correlation between educational attainment^58^ and SCDC (r_g_ = −0.30± 0.11, P = 0.007).

### Genomic inflation factor, heritability, and functional enrichment for the SQ-R GWAS

LDSR^91,92^ was used to calculate for inflation in test statistics due to unaccounted population stratification. Heritability was calculated using LDSR using the north-west European LD scores. Difference in heritability between males and females was quantified using:

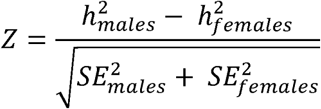

where Z is the Z score for the difference in heritability for a trait, (h^2^_males_ - h^2^_females_) is the difference h^2^_SNP_ estimate in males and females, and SE is the standard errors for heritability. Two-tailed P-values were calculated and reported as significant if P < 0.05.

For the primary GWAS (non-stratified analyses), we conducted functional annotation using FUMA^93^. We restricted our analyses to the non-stratified analyses due to the high genetic correlation between the sexes and the low statistical power of the sex-stratified GWAS. We conducted gene-based association analyses using MAGMA^94^ within FUMA and report significant genes after using a stringent Bonferroni corrected P < 0.05. In addition, we conducted enrichment for tissue specific expression and pathway analyses within FUMA. For the significant SNPs, we investigated enrichment for eQTLs using brain tissues in the BRAINEAC and GTEx^95^ database within FUMA. We further conducted partitioned heritability for tissue specific active chromatin marks and baseline functional categories using extended methods in LDSR^96^.

### Hi-C based annotations of fine mapped loci

We fine mapped three genome-wide significant loci (index SNPs: rs4146336 and rs1559586 or SQ; rs8005092 for SQ-R males) to obtain credible SNPs. First, we selected SNPs with P<0.01 that are located in the linkage disequilibrium (LD) region (r^2^>0.6) with an index SNP. LD structure within a locus was constructed by calculating correlations between SNPs within a locus (1KG v20130502). CAVIAR^97^ was then applied to the summary association statistics and LD structure for each index SNP to generate potentially causal (credible) SNPs with a posterior probability of 0.95. In total, we identified 14 credible SNPs from the three GWS loci.

For each locus, candidate genes were identified by mapping credible SNPs based on physical interactions in foetal brain as previously described^98^. One locus (index SNP rs4146336) was mapped to two genes, *LSAMP* and *PTMAP8*, indicating that two credible SNPs (rs13066948 and rs11713893) located in this locus physically interact with these genes.

### Genetic correlation

For all phenotypes we performed genetic correlation without constraining the intercept using LDSR. We identified significant genetic correlations using a Bonferroni adjusted P-value < 0.05. For the primary genetic correlation analysis with SQ-R, we included psychiatric conditions^57,63,99–102^, personality traits^103–105^, measures of intelligence^58,59,106,107^, and social traits related to autism^46,55^ including scores on the SCDC, as previous research has investigated the phenotypic correlation between these domains and systemizing^10,78,108–112^.

To understand the correlation between systemizing and various phenotypes that have been genetically correlated with autism, we used GWAS data from 15 phenotypes including autism. 10 of these phenotypes (cognitive aptitude^59^, educational attainment^58^, tiredness^113^, neuroticism^103^, subjective wellbeing^103^, schizophrenia^114^, major depression^102^, depressive symptoms^103^, ADHD^63^ and chronotype^115^), have been previously reported to be significantly genetically correlated with autism out of 234 phenotypes tested using LDHub^62^ (P < 2.1×10^−4^). We excluded college degree from this list, as previous work has identified near perfect genetic correlation between educational attainment and college dsegree^58^. In addition, we included data from friendship satisfaction^55^, family satisfaction^55^, systemizing, and self-reported empathy^46^, all of which are also significantly genetically correlated with autism with P < 2.1×10^−4^. These four additional phenotypes were not included in the previous paper which investigated genetic correlations with autism. Details of sample sizes with PMIDs/DOIs are provided in **Supplementary Table 12**. Cross trait genetic correlations were computed for all 15 phenotypes, and results were corrected for multiple testing using Bonferroni correction. A correlogram was created after using hierarchical clustering to cluster the phenotypes.

To investigate if the combination of negative genetic correlation social traits and positive genetic correlation for non-social traits is specific to autism, we conducted a genetic correlation between all psychiatric conditions for which we had access to summary GWAS statistics (ADHD^63^, Anxiety^116^, Autism^57^, Anorexia^101^, Bipolar Disorder^99^, Major Depressive Disorder^102^, OCD^117,118^, PTSD^119^, and Schizophrenia^114^) and SQ-R, self-reported empathy measured using the EQ^46^ and friendship satisfaction^55^. We chose friendship satisfaction and self-reported empathy as representative of social traits as these are the most relevant to the social domain of autism that we had access to GWAS summary statistics. The EQ is a short, 40-item self-report measure of empathy, which has been widely used and has good psychometric properties^60,120^. For instance, in the DSM-5, one of the criteria for autism is difficulties in making friends^1^. Additionally, differences in aspects of empathy compared to the neurotypical population have been widely reported in autism^50,51,121^, and is one of the items in measures such as ADOS-G.

### GWIS and GSEM

To investigate if the SQ-R is genetically correlated with autism independent of the genetic effects of educational attainment, we constructed a unique SQ-R phenotype after conditioning on the genetic effects of educational attainment using GWIS^122^. GWIS takes into account the genetic covariance between the two phenotypes to calculate the unique component of the phenotypes as a function of the genetic covariance and the h^2^. Prior to performing GWIS, we standardized the beta coefficients for the SQ-R GWAS by using the following formula:

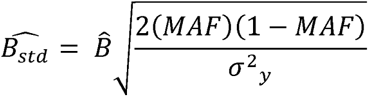

Where 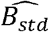 is the standardized regression coefficients, 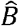 is the regression coefficient obtained from the non-standardized GWAS, MAF is the minor allele frequency, *σ*^2^ _*y*_ is the variance of the SQ-R. This equation is explained in detail in the **Supplementary Note**. We conducted GWIS using only educational attainment as we were unclear if the GWAS of cognitive aptitude^59^ was conducted on a standardized phenotype. Further, there is a high genetic correlation between cognitive aptitude and educational attainment. In addition to GWIS, to validate the findings, we conducted GSEM^123^, a complementary but independent method. GSEM uses the correlations and covariances calculated using LDSR after accounting for sample overlap.

### Polygenic scores in the SSC, AGRE, EU-AIMS LEAP and Paris cohorts

We generated polygenic scores for SQ-R (mean weighted score of all the alleles that contribute to higher systemizing) in 2,221 probands from the Simons Simplex Collection (Discovery dataset). We downloaded genotype data from the SSC from SFARI base (https://www.sfari.org/resource/sfari-base/). Individuals were genotyped on three different platforms: Illumina Omni2.5, Illumina 1Mv3, or Illumina 1Mv1. Informed consent or assent was obtained from all participants. In addition, the research team obtained ethical approval from the Cambridge Human Biology Research Ethics Committee to access and analyse the de-identified data from the Simons Simplex collection. We conducted a stringent quality control and imputation to generate genotypes used in this analysis for each of the platforms separately. The full pipeline is available here: https://github.com/autism-research-centre/SSC_liftover_imputation. Briefly, individuals were excluded if they had: a genotyping rate < 95%, excessive or low heterozygosity (less or more than 3 SD from the mean), mismatched reported and genetic sex, and families with mendelian errors > 5%. We further removed SNPs that significantly deviated from Hardy-Weinberg Equilibrium (P < 1×10^−6^), had mendelian errors in more than 10% of the families, and SNPs that were not genotyped in more than 10% of the families. We then conducted multidimensional scaling using the HapMap3 phase 3 population using the unrelated individuals CEU and TSI populations as representatives of the European population. This was conducted only in the parents to retain unrelated individuals for multidimensional scaling. Genetic principal components were calculated using only SNPs with minor allele frequency > 5%, and pruning the SNPs in Plink using an r^2^ of 0.2. We excluded families from further downstream analyses if either one the parents were greater or less than 5 standard deviations from the means of the first two genetic principal components calculated using only the unrelated individuals in HapMap3 CEU and TSI populations. Quality control was done using Plink v 1.9 and R. Phasing and imputation was conducted using the Michigan Imputation Server (https://imputationserver.sph.umich.edu/start.html) using the 1000 genomes Phase 3 v5 as the reference panel.

Polygenic scores were generated using PRSice2 (https://choishingwan.github.io/PRSice/) for the SQ-R using the non-stratified GWAS data. We calculated the mean polygenic score for each of the 2,221 probands in the SSC, after clumping SNPs using an R^2^ threshold of 0.1. Prior to generating polygenic scores, we confirmed that the probands were not related to each other using identity by descent PI-HAT > 0.15 as a relatedness cut-off. We used a P-value threshold of 1 as previous research on educational attainment, subjective wellbeing and social relationship satisfaction, all suggest that the maximum variance explained is at a threshold of 1^58,103^. This is expected for highly polygenic traits where many SNPs incrementally contribute to the variance explained^124^. Polygenic scoring was done using standardized scores on two different phenotypes as the dependent variable (RBS-R and the social and communication domain of the ADOS-G). We included sex, platform, the first 15 genetic principal components and standardized full-scale IQ as covariates. In addition, for the analysis of ADOS-G, we included the ADOS-G module as a covariate. Linear regression was conducted in R. A total of 135,233 SNPs were included in the polygenic score analyses after clumping and thresholding.

To validate the polygenic scores, we conducted additional polygenic score analysis using data combined from the AGRE, EU-AIMS LEAP and Paris cohorts. We followed similar quality control and imputation procedures to the SSC cohort. Given that this dataset was a mix of related and unrelated individuals, we chose unrelated individuals using a genomic relationship matrix (GRM) as provided in GCTA (--grm-cutoff 0.05)^125^. To calculate GRMs, we included only SNPs with minor allele frequency > 1%. Scripts are provided here: https://github.com/vwarrier/PARIS_LEAP_analysis. Polygenic scores were calculated using PRSice2 as described for the SSC data. Given the differences in dataset, polygenic scores were calculated separately for the AGRE dataset, and the EU-AIMS LEAP and Paris datasets combined. For each regression, we included sex and the first 10 genetic principal components (standardized). The dependent variables were standardized scores on the RBS-R (N = 426) and the ADOS-G social and communication subscale (N = 475). IQ information was unavailable for most individuals, and hence we did not include IQ as a covariate. We combined the results of the EU-AIMS LEAP and Paris cohorts, and the AGRE dataset using inverse variance weighted fixed-effect meta-analysis using the formula below:

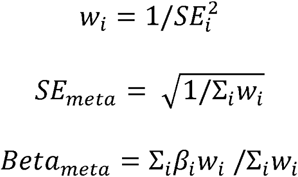

Where *β*_*i*_ is the standardized regression coefficient of the polygenic scores, *SE*_*i*_ is the associated standard error, and *W*_*i*_ is the weight.

### Bivariate GREML

We conducted bivariate genetic correlation using GCTA GREML to test the genetic correlation between the ADOS social and communication domains and the RBS-R scores. We created a GRM after including autistic individuals from the SSC, AGRE, EU-AIMS LEAP and Paris cohorts. We excluded SNPs and individuals using the same quality control pipeline as applied to the SSC dataset outlined in the section above. We further restricted our analysis only to SNPs with a minor allele frequency > 1%. We excluded related individuals (--grm-cutoff 0.05) resulting in a total of 2,989 individuals. Of this, 2,652 individuals had scores for the ADOS social and communication domain and 2,550 individuals had scores on the RBS-R. We included sex and the first 10 genetic principal components as covariates.

## Data availability

The SQ-R GWAS results are available from 23andMe. The full set of summary statistics can be made available to qualified investigators who enter into an agreement with 23andMe that protects participant confidentiality. Interested investigators should for more information. Top SNPs (n = 10,000) can be visualized here: https://ghfc.pasteur.fr. Data for ALSPAC can be requested here: http://www.bristol.ac.uk/alspac/researchers/access/. Data from the Simons Simplex Collection can be requested here: https://www.sfari.org/resource/sfari-base/. Summary GWAS statistics were downloaded from the PGC consortium: http://www.med.unc.edu/pgc/results-and-downloads. Data for chronotype was downloaded from http://www.t2diabetesgenes.org/data/. Data for self-reported tiredness was downloaded from http://www.ccace.ed.ac.uk/node/335.

## Software and code availability

Genomic-SEM: https://github.com/MichelNivard/GenomicSEM

GWIS: https://sites.google.com/site/mgnivard/gwis

Plink: https://www.cog-genomics.org/plink2/

PRSice2: https://choishingwan.github.io/PRSice/

CAVIAR: http://genetics.cs.ucla.edu/caviar/

Michigan Imputation Server: https://imputationserver.sph.umich.edu/index.html

Custom code for quality control of the SSC and the other cohorts can be downloaded from https://github.com/autism-research-centre/SSC_liftover_imputation (DOI: 10.5281/zenodo.3342561) and from https://github.com/vwarrier/PARIS_LEAP_analysis (DOI: 10.5281/zenodo.3342569)

## Acknowledgements

We are grateful to Michel Nivard and Beate St. Pourcain for their help with the analytical methods. We thank the research participants and employees of 23andMe for making this work possible. We are grateful to all the families who took part in this study, the midwives for their help in recruiting them, and the whole ALSPAC team, which includes interviewers, computer and laboratory technicians, clerical workers, research scientists, volunteers, managers, receptionists, and nurses. VW was funded by St. John’s College, Cambridge, and the Cambridge Commonwealth Trust. This study was funded by grants to SBC from the Medical Research Council, the Wellcome Trust, the Autism Research Trust, the Templeton World Charity Foundation,, and to TB from the Institut Pasteur, the CNRS, the INSERM, The Fondamental Foundation, the APHP, the BioPsy Labex and the University Paris Diderot. The research was conducted in association with the National Institute for Health Research (NIHR) Collaboration for Leadership in Applied Health Research and Care East of England at Cambridgeshire and Peterborough NHS Foundation Trust. We also received support from the NIHR Cambridge Biomedical Research Centre. We acknowledge with gratitude the generous support of Drs Dennis and Mireille Gillings in strengthening the collaboration between SBC and TB, and between Cambridge University and the Institut Pasteur. The views expressed are those of the author(s) and not necessarily those of the NHS, the NIHR or the Department of Health. Data obtained from 23andMe was supported by the National Human Genome Research Institute of the National Institutes of Health (grant number R44HG006981). The UK Medical Research Council and Wellcome (grant ref: 102215/2/13/2) and the University of Bristol provide core support for ALSPAC. GWAS data was generated by Sample Logistics and Genotyping Facilities at Wellcome Sanger Institute and LabCorp (Laboratory Corporation of America) using support from 23andMe. This publication is the work of the authors who will serve as guarantors for the content of this paper. The iPSYCH (The Lundbeck Foundation Initiative for Integrative Psychiatric Research) team acknowledges funding from The Lundbeck Foundation (grant no R102-A9118 and R155-2014-1724), the Stanley Medical Research Institute, the European Research Council (project no: 294838), the Novo Nordisk Foundation for supporting the Danish National Biobank resource, and grants from Aarhus and Copenhagen Universities and University Hospitals, including support to the iSEQ Center, the GenomeDK HPC facility, and the CIRRAU Center. The project leading to this application has received funding from the Innovative Medicines Initiative 2 Joint Undertaking (JU) under grant agreement No 777394. The JU receives support from the European Union’s Horizon 2020 research and innovation programme and EFPIA and AUTISM SPEAKS, Autistica, SFARI. We thank the iPSCH-Broad Autism Group and the EU-AIMS LEAP group for sharing data. A full list of the authors and affiliations in the iPSYCH-Broad autism group and the EU-AIMS LEAP group is provided in the Supplementary Information.

## Author contributions

VW, TB, SBC, and DAH conceived and designed the analysis. CSL, FC, RD, WDW, JB, ADB, JG, GP, 23andMe Research Team, and DAH collected or contributed the data or analysis tools. VW, RT, HW, FC, and WDW performed the analysis. VW, RT, BC, DAH, TB, and SBC wrote the paper. DAH, TB, and SBC supervised the analysis.

## †23andMe Research Team

Michelle Agee^1^, Babak Alipanahi^1^, Adam Auton^1^, Robert K. Bell^1^, Katarzyna Bryc^1^, Sarah L. Elson^1^, Pierre Fontanillas^1^, Nicholas A. Furlotte^1^, Karen E. Huber^1^, Aaron Kleinman^1^, Nadia K. Litterman^1^, Jennifer C. McCreight^1^, Matthew H. McIntyre^1^, Joanna L. Mountain^1^, Carrie A.M. Northover^1^, Steven J. Pitts^1^, J. Fah Sathirapongsasuti^1^, Olga V. Sazonova^1^, Janie F. Shelton^1^, Suyash Shringarpure^1^, Chao Tian^1^, Joyce Y. Tung^1^, Vladimir Vacic^1^, and Catherine H. Wilson^1^.

^17^ 23andMe Inc., Mountain View, California 94043, USA

## References

1. American Psychiatric Association. Diagnostic and statistical manual of mental disorders (5th ed.). (2013).

2. Happé, F. & Ronald, A. The ‘Fractionable Autism Triad’: A Review of Evidence from Behavioural, Genetic, Cognitive and Neural Research. Neuropsychol. Rev. 18, 287–304 (2008).

3. Happé, F., Ronald, A. & Plomin, R. Time to give up on a single explanation for autism. Nat. Neurosci. 9, 1218–1220 (2006).

4. Shuster, J., Perry, A., Bebko, J. & Toplak, M. E. Review of Factor Analytic Studies Examining Symptoms of Autism Spectrum Disorders. J. Autism Dev. Disord. 44, 90–110 (2014).

5. Mandy, W. P. L. & Skuse, D. H. Research Review: What is the association between the social-communication element of autism and repetitive interests, behaviours and activities? J. Child Psychol. Psychiatry 49, 795–808 (2008).

6. Palmer, C. J., Paton, B., Enticott, P. G. & Hohwy, J. ‘Subtypes’ in the Presentation of Autistic Traits in the General Adult Population. J. Autism Dev. Disord. 45, 1291–1301 (2015).

7. Guthrie, W., Swineford, L. B., Wetherby, A. M. & Lord, C. Comparison of DSM-IV and DSM-5 Factor Structure Models for Toddlers With Autism Spectrum Disorder. J. Am. Acad. Child Adolesc. Psychiatry 52, 797–805.e2 (2013).

8. Frazier, T. W., Youngstrom, E. A., Kubu, C. S., Sinclair, L. & Rezai, A. Exploratory and Confirmatory Factor Analysis of the Autism Diagnostic Interview-Revised. J. Autism Dev. Disord. 38, 474–480 (2008).

9. Grove, R., Baillie, A., Allison, C., Baron-Cohen, S. & Hoekstra, R. A. Empathizing, systemizing, and autistic traits: Latent structure in individuals with autism, their parents, and general population controls. J. Abnorm. Psychol. 122, 600–609 (2013).

10. Wheelwright, S. J. et al. Predicting Autism Spectrum Quotient (AQ) from the Systemizing Quotient-Revised (SQ-R) and Empathy Quotient (EQ). Brain Res. 1079, 47–56 (2006).

11. Svedholm-Häkkinen, A. M., Halme, S. & Lindeman, M. Empathizing and systemizing are differentially related to dimensions of autistic traits in the general population. Int. J. Clin. Heal. Psychol. 18, 35–42 (2018).

12. Greenberg, D. M., Warrier, V., Allison, C. & Baron-Cohen, S. Testing the Empathizing-Systemizing theory of sex differences and the Extreme Male Brain theory of autism in half a million people. Proc. Natl. Acad. Sci. U. S. A. 201811032 (2018). doi:10.1073/pnas.1811032115

13. Ronald, A., Happe, F. & Plomin, R. The genetic relationship between individual differences in social and nonsocial behaviours characteristic of autism. Dev. Sci. 8, 444–458 (2005).

14. Ronald, A. et al. Genetic Heterogeneity Between the Three Components of the Autism Spectrum: A Twin Study. J. Am. Acad. Child Adolesc. Psychiatry 45, 691–699 (2006).

15. Dworzynski, K., Happé, F., Bolton, P. & Ronald, A. Relationship Between Symptom Domains in Autism Spectrum Disorders: A Population Based Twin Study. J. Autism Dev. Disord. 39, 1197–1210 (2009).

16. Norbury, C. F. Practitioner Review: Social (pragmatic) communication disorder conceptualization, evidence and clinical implications. J. Child Psychol. Psychiatry 55, 204–216 (2014).

17. Uljarević, M., Evans, D. W., Alvares, G. A. & Whitehouse, A. J. O. Short report: relationship between restricted and repetitive behaviours in children with autism spectrum disorder and their parents. Mol. Autism 7, 29 (2016).

18. Georgiades, S. et al. Investigating phenotypic heterogeneity in children with autism spectrum disorder: a factor mixture modeling approach. J. Child Psychol. Psychiatry 54, 206–215 (2013).

19. St Pourcain, B. et al. Variability in the common genetic architecture of social-communication spectrum phenotypes during childhood and adolescence. Mol. Autism 5, 18 (2014).

20. Cantor, R. M. et al. ASD restricted and repetitive behaviors associated at 17q21.33: genes prioritized by expression in fetal brains. Mol. Psychiatry (2017). doi:10.1038/mp.2017.114

21. Alarcón, M., Cantor, R. M., Liu, J., Gilliam, T. C. & Geschwind, D. H. Evidence for a language quantitative trait locus on chromosome 7q in multiplex autism families. Am. J. Hum. Genet. 70, 60–71 (2002).

22. Lowe, J. K., Werling, D. M., Constantino, J. N., Cantor, R. M. & Geschwind, D. H. Social responsiveness, an autism endophenotype: genomewide significant linkage to two regions on chromosome 8. Am. J. Psychiatry 172, 266–75 (2015).

23. Shao, Y. et al. Fine mapping of autistic disorder to chromosome 15q11-q13 by use of phenotypic subtypes. Am. J. Hum. Genet. 72, 539–48 (2003).

24. Cannon, D. S. et al. Genome-wide linkage analyses of two repetitive behavior phenotypes in Utah pedigrees with autism spectrum disorders. Mol. Autism 1, 3 (2010).

25. Tao, Y. et al. Evidence for contribution of common genetic variants within chromosome 8p21.2-8p21.1 to restricted and repetitive behaviors in autism spectrum disorders. BMC Genomics 17, 163 (2016).

26. Ruzich, E. et al. Measuring autistic traits in the general population: a systematic review of the Autism-Spectrum Quotient (AQ) in a nonclinical population sample of 6,900 typical adult males and females. Mol. Autism 6, 2 (2015).

27. Baron-Cohen, S., Wheelwright, S. J., Skinner, R., Martin, J. & Clubley, E. The autism-spectrum quotient (AQ): evidence from Asperger syndrome/high-functioning autism, males and females, scientists and mathematicians. J. Autism Dev. Disord. 31, 5–17 (2001).

28. Posserud, M.-B., Lundervold, A. J. & Gillberg, C. Autistic features in a total population of 7-9-year-old children assessed by the ASSQ (Autism Spectrum Screening Questionnaire). J. Child Psychol. Psychiatry 47, 167–175 (2006).

29. Constantino, J. N. & Todd, R. D. Autistic Traits in the General Population. Arch. Gen. Psychiatry 60, 524 (2003).

30. Constantino, J. N. et al. The factor structure of autistic traits. J. Child Psychol. Psychiatry 45, 719–726 (2004).

31. Hoekstra, R. A. et al. Heritability of autistic traits in the general population. Arch. Pediatr. Adolesc. Med. 161, 372–7 (2007).

32. Ronald, A. & Hoekstra, R. A. Autism spectrum disorders and autistic traits: A decade of new twin studies. Am. J. Med. Genet. Part B Neuropsychiatr. Genet. 156, 255–274 (2011).

33. de Zeeuw, E. L., van Beijsterveldt, C. E. M., Hoekstra, R. A., Bartels, M. & Boomsma, D. I. The etiology of autistic traits in preschoolers: a population-based twin study. J. Child Psychol. Psychiatry 58, 893–901 (2017).

34. Wheelwright, S. J., Auyeung, B., Allison, C. & Baron-Cohen, S. Defining the broader, medium and narrow autism phenotype among parents using the Autism Spectrum Quotient (AQ). Mol. Autism 1, 10 (2010).

35. Sucksmith, E., Roth, I. & Hoekstra, R. A. Autistic Traits Below the Clinical Threshold: Re-examining the Broader Autism Phenotype in the 21st Century. Neuropsychol. Rev. 21, 360–389 (2011).

36. Sasson, N. J., Lam, K. S., Parlier, M., Daniels, J. L. & Piven, J. Autism and the broad autism phenotype: familial patterns and intergenerational transmission. J. Neurodev. Disor.. 5, 11 (2013).

37. Constantino, J. N. & Todd, R. D. Intergenerational transmission of subthreshold autistic traits in the general population. Biol. Psychiatry 57, 655–660 (2005).

38. Murray, A. L., McKenzie, K., Kuenssberg, R. & Booth, T. Do the Autism Spectrum Quotient (AQ) and Autism Spectrum Quotient Short Form (AQ-S) Primarily Reflect General ASD Traits or Specific ASD Traits? A Bi-Factor Analysis. Assessment 24, 444–457 (2017).

39. Lundström, S. et al. Autism Spectrum Disorders and Autisticlike Traits. Arch. Gen. Psychiatry 69, 46 (2012).

40. Robinson, E. B. et al. Evidence that autistic traits show the same etiology in the general population and at the quantitative extremes (5%, 2.5%, and 1%). Arch. Gen. Psychiatry 68, 1113–21 (2011).

41. Ronald, A., Happé, F., Price, T. S., Baron-Cohen, S. & Plomin, R. Phenotypic and genetic overlap between autistic traits at the extremes of the general population. J. Am. Acad. Child Adolesc. Psychiatr 45, 1206–14 (2006).

42. Colvert, E. et al. Heritability of autism spectrum disorder in a UK population-based twin sample. JAMA Psychiatry 72, 415–23 (2015).

43. Bralten, J. et al. Autism spectrum disorders and autistic traits share genetics and biology. Mol. Psychiatry (2017). doi:10.1038/mp.2017.98

44. St Pourcain, B. et al. ASD and schizophrenia show distinct developmental profiles in common genetic overlap with population-based social communication difficulties. Mol. Psychiatry (2017). doi:10.1038/mp.2016.198

45. Robinson, E. B. et al. Genetic risk for autism spectrum disorders and neuropsychiatric variation in the general population. Nat. Genet. 48, 552–5 (2016).

46. Warrier, V. et al. Genome-wide analyses of self-reported empathy: Correlations with autism, schizophrenia, and anorexia nervosa. Transl. Psychiatry 8, (2018).

47. Warrier, V. et al. Genome-wide meta-analysis of cognitive empathy: heritability, and correlates with sex, neuropsychiatric conditions and cognition. Mol. Psychiatry (2017).

48. Cantor, R. M. et al. ASD restricted and repetitive behaviors associated at 17q21.33: genes prioritized by expression in fetal brains. Mol. Psychiatry 23, 993–1000 (2018).

49. Baron-Cohen, S. The hyper-systemizing, assortative mating theory of autism. Prog. Neuropsychopharmacol. Biol. Psychiatry 30, 865–72 (2006).

50. Auyeung, B. et al. The Children’s Empathy Quotient and Systemizing Quotient: Sex Differences in Typical Development and in Autism Spectrum Conditions. J. Autism Dev. Disord. 39, 1509–1521 (2009).

51. Baron-Cohen, S. et al. Attenuation of typical sex differences in 800 adults with autism vs. 3,900 controls. PLoS One 9, e102251 (2014).

52. Asperger, H. ‘Autistic psychopathy’ in childhood. in Autism and Asperger syndrome (ed. Frith, U.) 37–92 (Cambridge University Press, 1944). doi:10.1017/CBO9780511526770.002

53. Kanner, L. Autistic disturbances of affective contact. Nerv. Child J. Psychopathol. Psychother. Ment. Hyg. Guid. Child 2 217–50 (1943).

54. Auyeung, B. et al. Foetal testosterone and the child systemizing quotient. Eur. J. Endocrinol. 155, S123–S130 (2006).

55. Warrier, V., the 23andMe Research team, Bourgeron, T. & Baron-Cohen, S. Genome-wide association study of social relationship satisfaction: significant loci and correlations with psychiatric conditions. bioRxiv 196071 (2017). doi:10.1101/196071

56. Palmer, E. E. et al. A Recurrent De Novo Nonsense Variant in ZSWIM6 Results in Severe Intellectual Disability without Frontonasal or Limb Malformations. Am. J. Hum. Genet. 101, 995–1005 (2017).

57. Grove, J. et al. Identification of common genetic risk variants for autism spectrum disorder. Nat. Genet. 51, 431–444 (2019).

58. Okbay, A. et al. Genome-wide association study identifies 74 loci associated with educational attainment. Nature 533, 539–542 (2016).

59. Sniekers, S. et al. Genome-wide association meta-analysis of 78,308 individuals identifies new loci and genes influencing human intelligence. Nat. Genet. (2017). doi:10.1038/ng.3869

60. Baron-Cohen, S. & Wheelwright, S. J. The Empathy Quotient: an investigation of adults with Asperger syndrome or high functioning autism, and normal sex differences. J. Autism Dev. Disord. 34, 163–75 (2004).

61. Skuse, D. H., Mandy, W. P. L. & Scourfield, J. Measuring autistic traits: heritability, reliability and validity of the Social and Communication Disorders Checklist. Br. J. Psychiatry 187, 568–572 (2005).

62. Zheng, J. et al. LD Hub: a centralized database and web interface to perform LD score regression that maximizes the potential of summary level GWAS data for SNP heritability and genetic correlation analysis. Bioinformatics 33, 272–279 (2016).

63. Demontis, D. et al. Discovery of the first genome-wide significant risk loci for attention deficit/hyperactivity disorder. Nat. Genet. 51, 63–75 (2019).

64. Ripke, S. et al. Genome-wide association analysis identifies 13 new risk loci for schizophrenia. Nat. Genet. 45, 1150–1159 (2013).

65. Stahl, E. et al. Genomewide association study identifies 30 loci associated with bipolar disorder. bioRxiv 173062 (2018). doi:10.1101/173062

66. Baron-Cohen, S. The extreme male brain theory of autism. Trends Cogn. Sci. 6, 248–254 (2002).

67. Tung, J. Y. et al. Efficient replication of over 180 genetic associations with self-reported medical data. PLoS One 6, e23473 (2011).

68. Do, C. B. et al. Web-based genome-wide association study identifies two novel loci and a substantial genetic component for Parkinson’s disease. PLoS Genet. 7, e1002141 (2011).

69. Eriksson, N. et al. Novel associations for hypothyroidism include known autoimmune risk loci. PLoS One 7, e34442 (2012).

70. Henn, B. M. et al. Cryptic distant relatives are common in both isolated and cosmopolitan genetic samples. PLoS One 7, e34267 (2012).

71. Boyd, A. et al. Cohort Profile: The ‘Children of the 90s’—the index offspring of the Avon Longitudinal Study of Parents and Children. Int. J. Epidemiol. 42, 111–127 (2013).

72. Fischbach, G. D. & Lord, C. The Simons Simplex Collection: A Resource for Identification of Autism Genetic Risk Factors. Neuron 68, 192–195 (2010).

73. Geschwind, D. H. et al. The autism genetic resource exchange: a resource for the study of autism and related neuropsychiatric conditions. Am. J. Hum. Genet. 69, 463–6 (2001).

74. Charman, T. et al. The EU-AIMS Longitudinal European Autism Project (LEAP): clinical characterisation. Mol. Autism 8, 27 (2017).

75. Durand, C. M. et al. Mutations in the gene encoding the synaptic scaffolding protein SHANK3 are associated with autism spectrum disorders. Nat. Genet. 39, 25–27 (2007).

76. Groen, Y., Fuermaier, A. B. M., Den Heijer, A. E., Tucha, O. & Althaus, M. The Empathy and Systemizing Quotient: The psychometric properties of the Dutch version and a review of the cross-cultural stability. J. Autism Dev. Disord. 45, 2848–64 (2015).

77. Allison, C., Baron-Cohen, S., Stone, M. H. & Muncer, S. J. Rasch modeling and confirmatory factor analysis of the systemizing quotient-revised (SQ-R) scale. Span. J. Psychol. 18, E16 (2015).

78. Ling, J., Burton, T. C., Salt, J. L. & Muncer, S. J. Psychometric analysis of the systemizing quotient (SQ) scale. Br. J. Psychol. 100, 539–552 (2009).

79. Wakabayashi, A. et al. Empathizing and systemizing in adults with and without autism spectrum conditions: Cross-cultural stability. J. Autism Dev. Disord. 37, 1823–1832 (2007).

80. Bölte, S., Westerwald, E., Holtmann, M., Freitag, C. M. & Poustka, F. Autistic traits and autism spectrum disorders: the clinical validity of two measures presuming a continuum of social communication skills. J. Autism Dev. Disord. 41, 66–72 (2011).

81. Lord, C. et al. The Autism Diagnostic Observation Schedule—Generic: A Standard Measure of Social and Communication Deficits Associated with the Spectrum of Autism. J. Autism Dev. Disord. 30, 205–223 (2000).

82. Gotham, K., Risi, S., Pickles, A. & Lord, C. The Autism Diagnostic Observation Schedule: Revised Algorithms for Improved Diagnostic Validity. J. Autism Dev. Disord. 37, 613–627 (2007).

83. Bodfish, J. W., Symons, F. J., Parker, D. E. & Lewis, M. H. Varieties of Repetitive Behavior in Autism: Comparisons to Mental Retardation. J. Autism Dev. Disord. 30, 237–243 (2000).

84. Lam, K. S. L. & Aman, M. G. The Repetitive Behavior Scale-Revised: Independent Validation in Individuals with Autism Spectrum Disorders. J. Autism Dev. Disord. 37, 855–866 (2007).

85. Browning, S. R. & Browning, B. L. Rapid and accurate haplotype phasing and missing-data inference for whole-genome association studies By use of localized haplotype clustering. Am. J. Hum. Genet. 81, 1084–1097 (2007).

86. Fuchsberger, C., Abecasis, G. R. & Hinds, D. A. minimac2: faster genotype imputation. Bioinformatics 31, 782–784 (2015).

87. Bigdeli, T. B. et al. A simple yet accurate correction for winner’s curse can predict signals discovered in much larger genome scans. Bioinformatics 32, 2598–2603 (2016).

88. Delaneau, O., Marchini, J. & Zagury, J.-F. A linear complexity phasing method for thousands of genomes. Nat. Methods 9, 179–181 (2011).

89. Howie, B. N., Donnelly, P. & Marchini, J. A flexible and accurate genotype imputation method for the next generation of genome-wide association studies. PLoS Genet. 5, e1000529 (2009).

90. Purcell, S. et al. PLINK: a tool set for whole-genome association and population-based linkage analyses. Am. J. Hum. Genet. 81, 559–75 (2007).

91. Bulik-Sullivan, B. K. et al. LD Score regression distinguishes confounding from polygenicity in genome-wide association studies. Nat. Genet. 47, 291–295 (2015).

92. Bulik-Sullivan, B. K. et al. An atlas of genetic correlations across human diseases and traits. Nat. Genet. 47, 1236–41 (2015).

93. Watanabe, K., Taskesen, E., van Bochoven, A. & Posthuma, D. Functional mapping and annotation of genetic associations with FUMA. Nat. Commun. 8, 1826 (2017).

94. de Leeuw, C. A., Mooij, J. M., Heskes, T. & Posthuma, D. MAGMA: generalized gene-set analysis of GWAS data. PLoS Comput. Biol. 11, 1–19 (2015).

95. Ardlie, K. G. et al. The Genotype-Tissue Expression (GTEx) pilot analysis: multitissue gene regulation in humans. Science (80-.). 348, 648–60 (2015).

96. Finucane, H. K. et al. Partitioning heritability by functional annotation using genome-wide association summary statistics. Nat. Genet. 47, 1228–1235 (2015).

97. Hormozdiari, F., Kostem, E., Kang, E. Y., Pasaniuc, B. & Eskin, E. Identifying Causal Variants at Loci with Multiple Signals of Association. Genetics 198, 497–508 (2014).

98. Won, H. et al. Chromosome conformation elucidates regulatory relationships in developing human brain. Nature 538, 523–527 (2016).

99. Sklar, P. et al. Large-scale genome-wide association analysis of bipolar disorder identifies a new susceptibility locus near ODZ4. Nat. Genet. 43, 977–983 (2011).

100. Pardiñas, A. F. et al. Common schizophrenia alleles are enriched in mutation-intolerant genes and in regions under strong background selection. Nat. Genet. 1 (2018). doi:10.1038/s41588-018-0059-2

101. Duncan, L. E. et al. Significant locus and metabolic genetic correlations revealed in genome-wide association study of anorexia nervosa. Am. J. Psychiatry appi.ajp.2017.1 (2017). doi:10.1176/appi.ajp.2017.16121402

102. Wray, N. R. et al. Genome-wide association analyses identify 44 risk variants and refine the genetic architecture of major depression. Nat. Genet. 50, 668–681 (2018).

103. Okbay, A. et al. Genetic variants associated with subjective well-being, depressive symptoms, and neuroticism identified through genome-wide analyses. Nat. Genet. 48, 624–633 (2016).

104. de Moor, M. H. M. et al. Meta-analysis of genome-wide association studies for personality. Mol. Psychiatry 17, 337–49 (2012).

105. van den Berg, S. M. et al. Meta-analysis of genome-wide association studies for extraversion: findings from the genetics of personality consortium. Behav. Genet. 46, 170–82 (2016).

106. Rietveld, C. A. et al. Common genetic variants associated with cognitive performance identified using the proxy-phenotype method. Proc. Natl. Acad. Sci. 111, 13790–13794 (2014).

107. Rietveld, C. A. et al. GWAS of 126,559 individuals identifies genetic variants associated with educational attainment. Science 340, 1467–71 (2013).

108. Hambrook, D., Tchanturia, K., Schmidt, U., Russell, T. & Treasure, J. Empathy, systemizing, and autistic traits in anorexia nervosa: a pilot study. Br. J. Clin. Psychol. 47, 335–9 (2008).

109. Russell-Smith, S. N., Bayliss, D. M., Maybery, M. T. & Tomkinson, R. L. Are the autism and positive schizotypy spectra diametrically opposed in empathizing and systemizing. J. Autism Dev. Disord. 43, 695–706 (2013).

110. Courty, A. et al. Levels of autistic traits in anorexia nervosa: a comparative psychometric study. BMC Psychiatry 13, 222 (2013).

111. Nettle, D. Empathizing and systemizing: what are they, and what do they contribute to our understanding of psychological sex differences? Br. J. Psychol. 98, 237–55 (2007).

112. Wakabayashi, A. & Kawashima, H. Is empathizing in the E–S theory similar to agreeableness? The relationship between the EQ and SQ and major personality domains. Pers. Individ. Dif. 76, 88–93 (2015).

113. Deary, V. et al. Genetic contributions to self-reported tiredness. Mol. Psychiatry 23, 609–620 (2018).

114. Ripke, S. et al. Biological insights from 108 schizophrenia-associated genetic loci. Nature 511, 421–7 (2014).

115. Jones, S. E. et al. Genome-Wide Association Analyses in 128,266 Individuals Identifies New Morningness and Sleep Duration Loci. PLOS Genet. 12, e1006125 (2016).

116. Otowa, T. et al. Meta-analysis of genome-wide association studies of anxiety disorders. Mol. Psychiatry (2016). doi:10.1038/mp.2015.197

117. Mattheisen, M. et al. Genome-wide association study in obsessive-compulsive disorder: results from the OCGAS. Mol. Psychiatry 20, 337–344 (2015).

118. International Obsessive Compulsive Disorder Foundation Genetics Collaborative (IOCDF-GC) and OCD Collaborative Genetics Association Studies (OCGAS), P. D. et al. Revealing the complex genetic architecture of obsessive-compulsive disorder using meta-analysis. Mol. Psychiatry 23, 1181–1188 (2018).

119. Duncan, L. E. et al. Largest GWAS of PTSD (N=20□070) yields genetic overlap with schizophrenia and sex differences in heritability. Mol. Psychiatry 23, 666–673 (2017).

120. Allison, C., Baron-Cohen, S., Wheelwright, S. J., Stone, M. H. & Muncer, S. J. Psychometric analysis of the Empathy Quotient (EQ). Pers. Individ. Dif. 51, 829–835 (2011).

121. Kok, F. M., Groen, Y., Becke, M., Fuermaier, A. B. M. & Tucha, O. Self-Reported Empathy in Adult Women with Autism Spectrum Disorders – A Systematic Mini Review. PLoS One 11, e0151568 (2016).

122. Nieuwboer, H. A., Pool, R., Dolan, C. V., Boomsma, D. I. & Nivard, M. G. GWIS: Genome-Wide Inferred Statistics for functions of multiple phenotypes. Am. J. Hum. Genet. 99, 917–927 (2016).

123. Grotzinger, A. D. et al. Genomic structural equation modelling provides insights into the multivariate genetic architecture of complex traits. Nat. Hum. Behav. 1 (2019). doi:10.1038/s41562-019-0566-x

124. Boyle, E. A., Li, Y. I. & Pritchard, J. K. An Expanded View of Complex Traits: From Polygenic to Omnigenic. Cell 169, 1177–1186 (2017).

125. Yang, J., Lee, S. H., Goddard, M. E. & Visscher, P. M. GCTA: a tool for genome-wide complex trait analysis. Am. J. Hum. Genet. 88, 76–82 (2011).

