## Supplementary Note for "Social and non-social autism symptom and trait domains are genetically dissociable"

### Supplementary Figure 1: Mean scores and standard deviations of the SQ-R

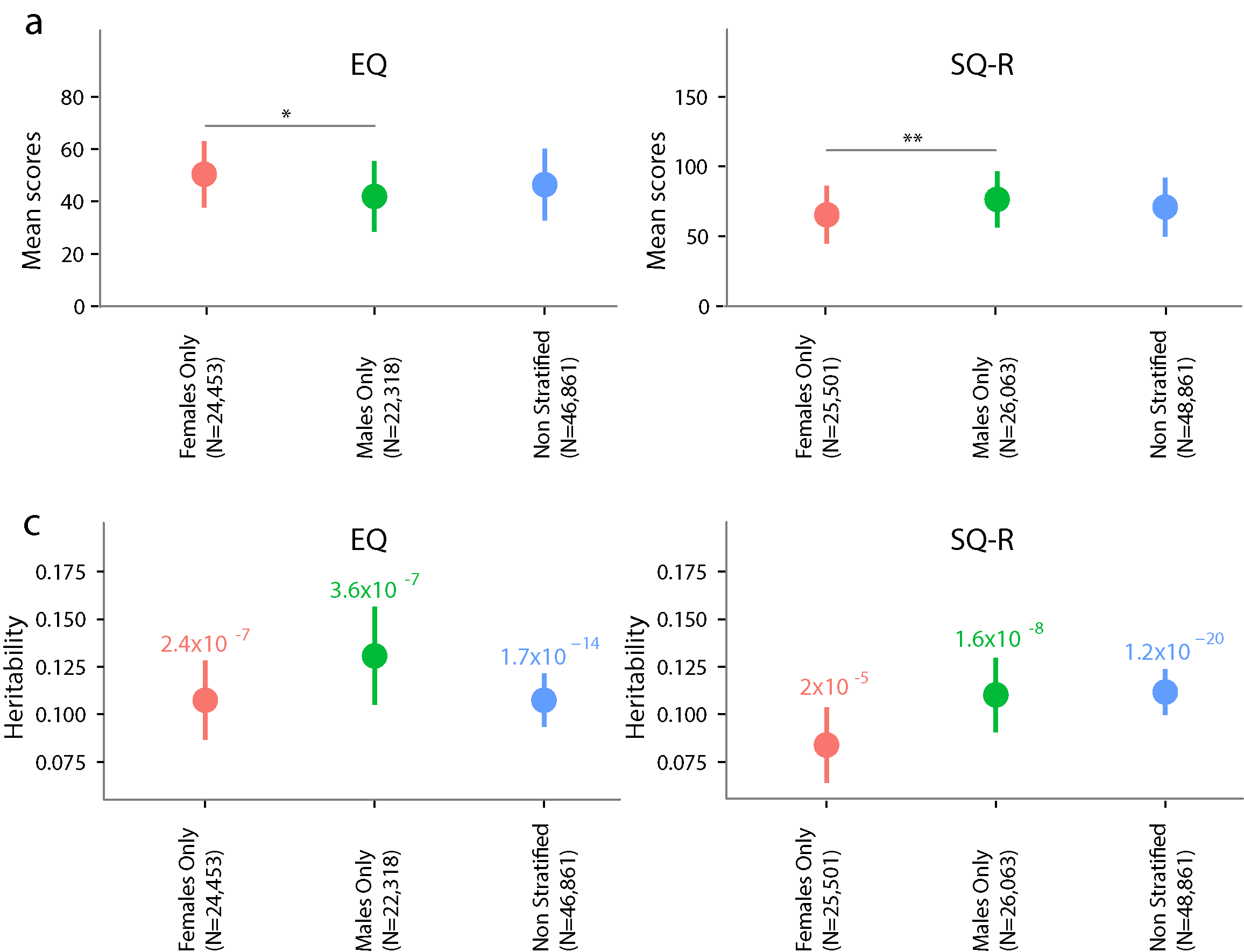

*Mean scores and standard deviations for the SQ-R scores for the three datasets. There was a significant difference in mean scores between males and females (P < 0.001, Cohen’s d = 0.54). Females: 65.4(20.6), Males: 76.5(20), Non-stratified: 71(21). Total score ranges from 0 – 150. Error bars represent the standard deviations.*

### Supplementary Figure 2: Regional LD plot for the three significant SNPs

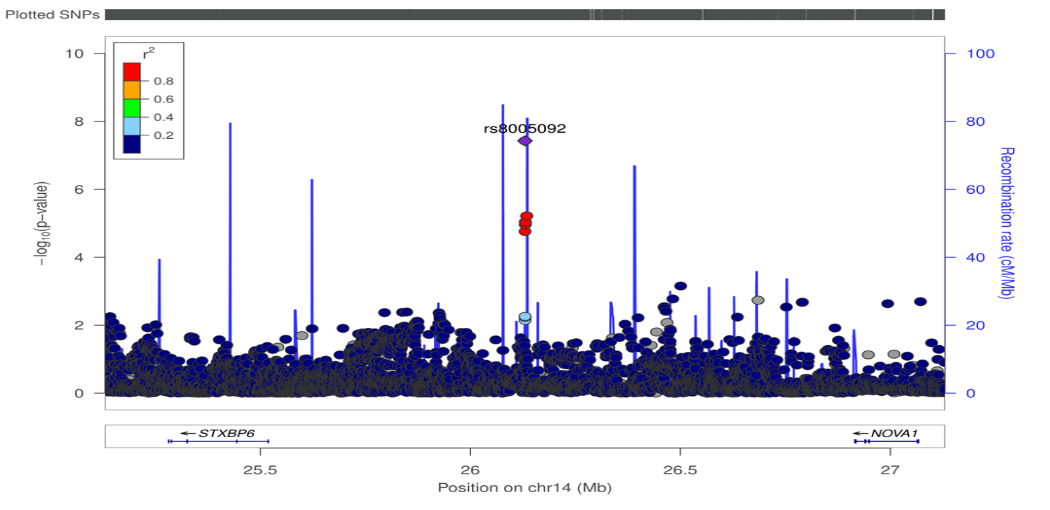

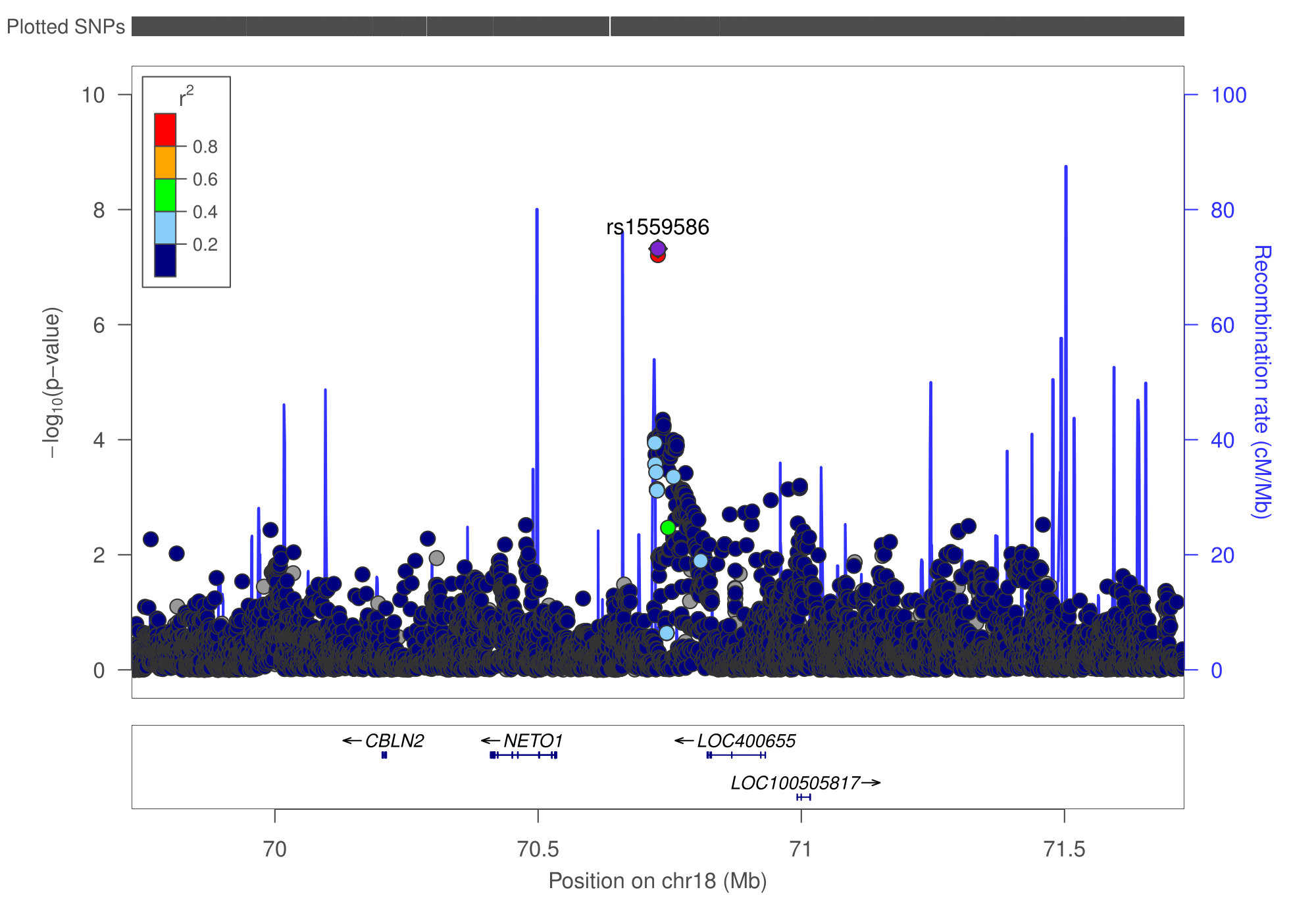

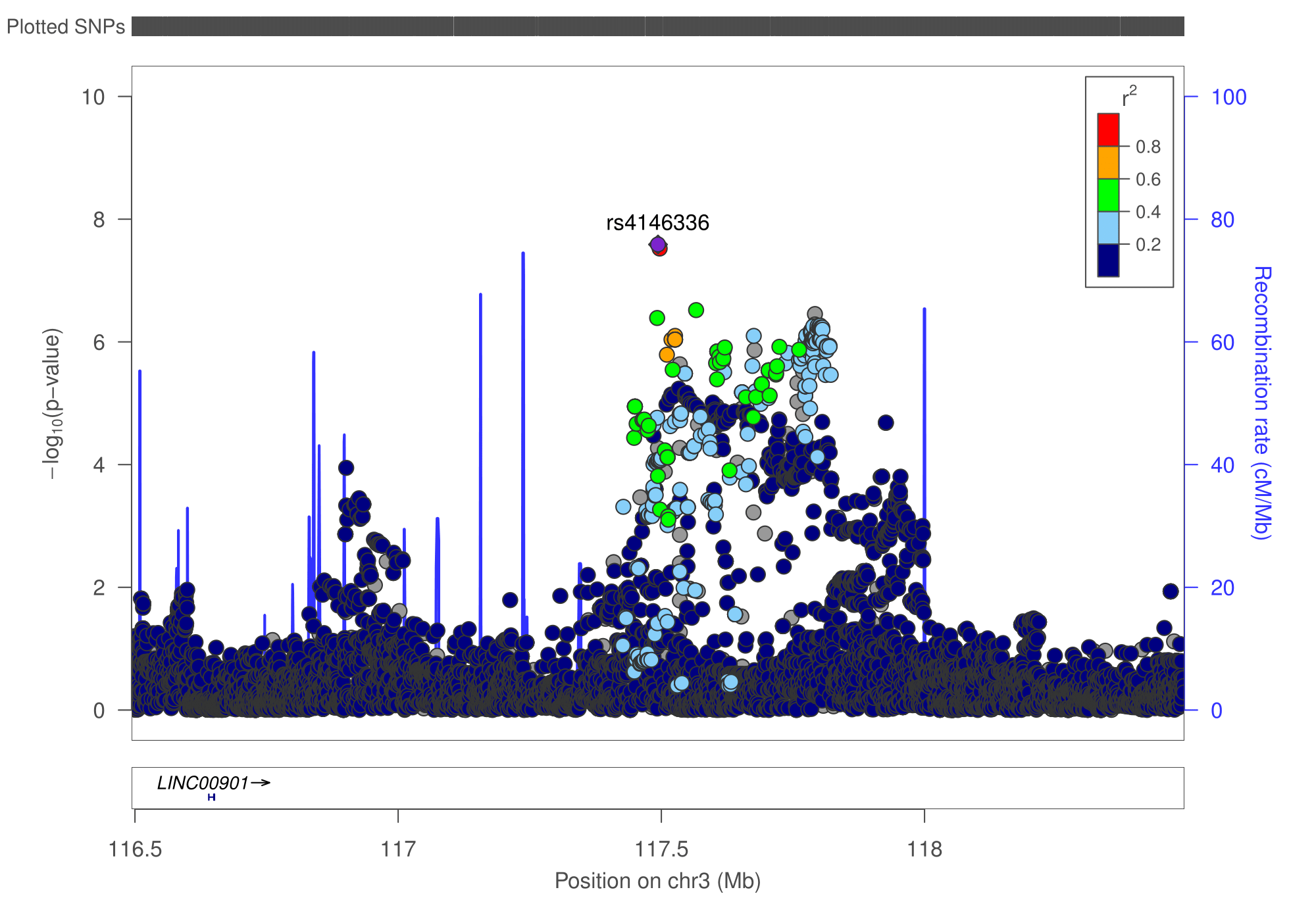

*Regional LD plots for the three significant SNPs. The top locus zoom plot is for rs8005092 in the males-only GWAS. The bottom two panels provide the regional LD plots for rs1559586 (left) and rs4146336 (right) in the non-stratified GWAS.*

### Supplementary Figure 3: Additive heritability for the three GWAS

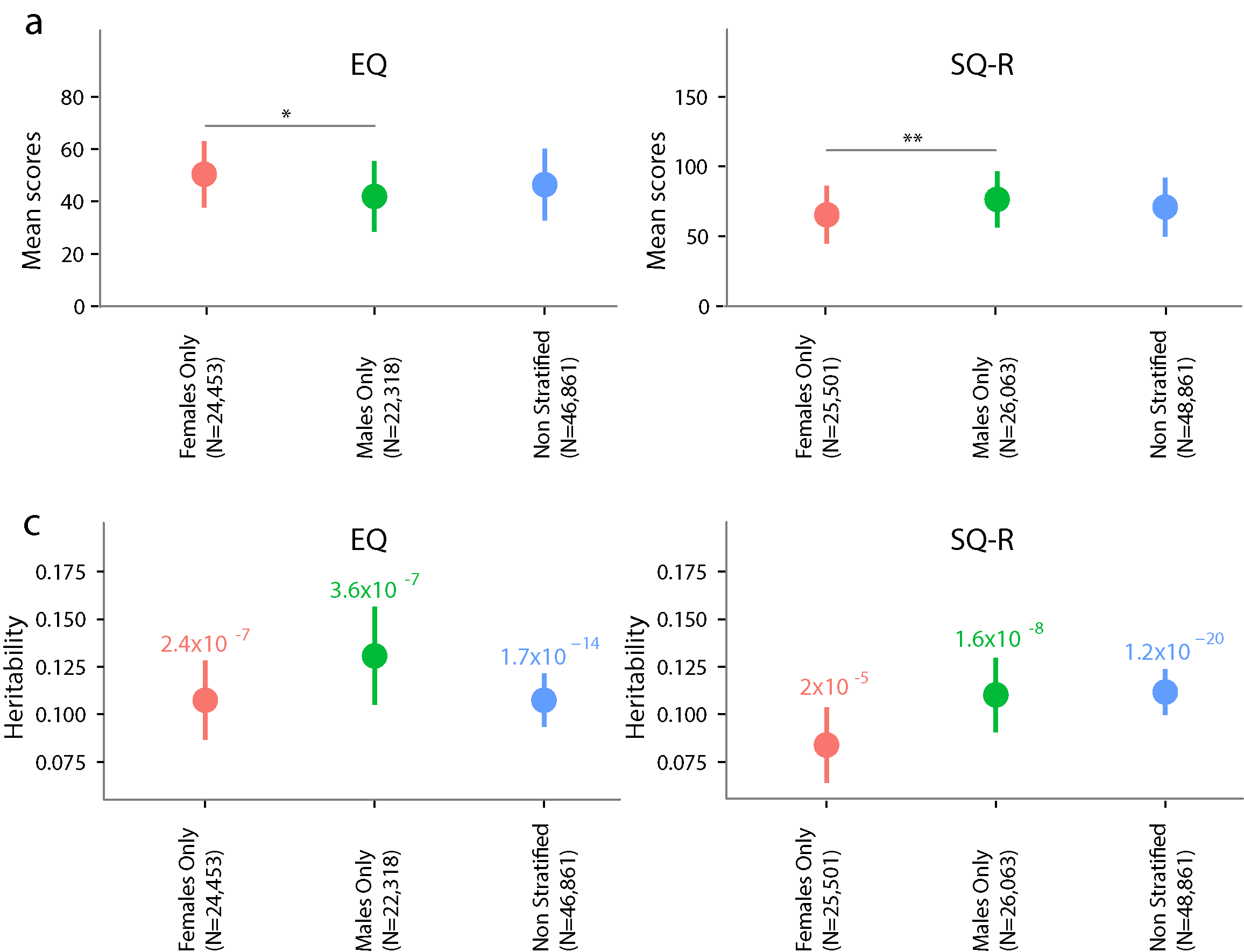

*Additive SNP heritability estimates for the SQ-R GWAS. Estimates provided for the non-stratified and the sex-stratified GWAS datasets. Error bars represent standard errors.*

### Supplementary Figure 4: Enrichment with active chromatin marks

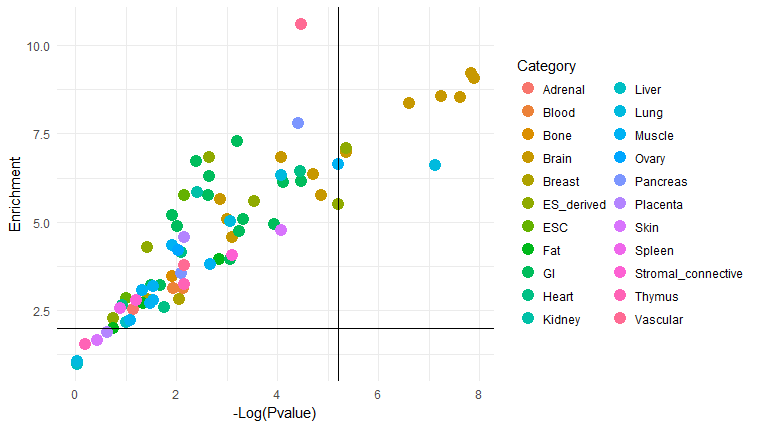

*The figure provides enrichment of active chromatin marks in the SQ-R GWAS based on category. The vertical line is the FDR-corrected P-value, and the horizontal line represents an enrichment of 2. The bulk of the significantly enriched categories are from the brain. Enrichment and P-values are provided in Supplementary Table 6.*

### Supplementary Figure 5: Tissue specific heritability

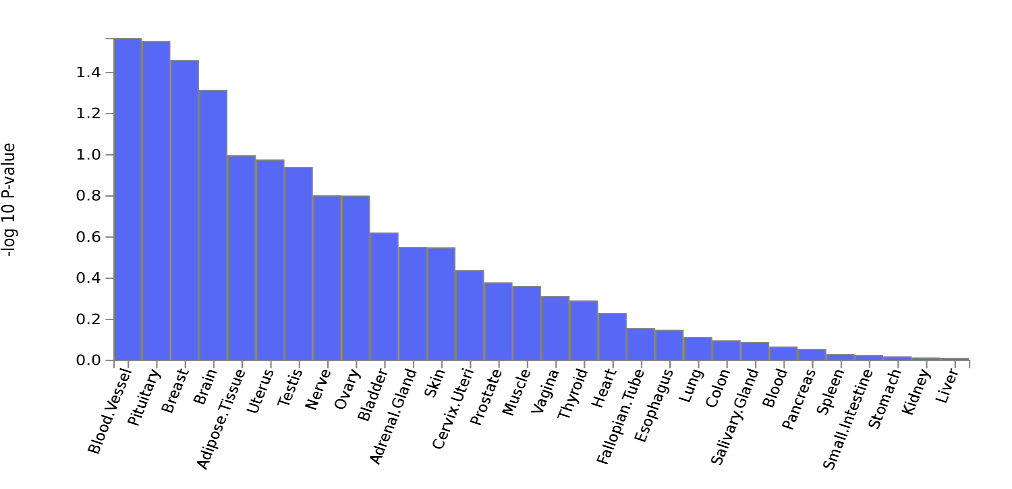

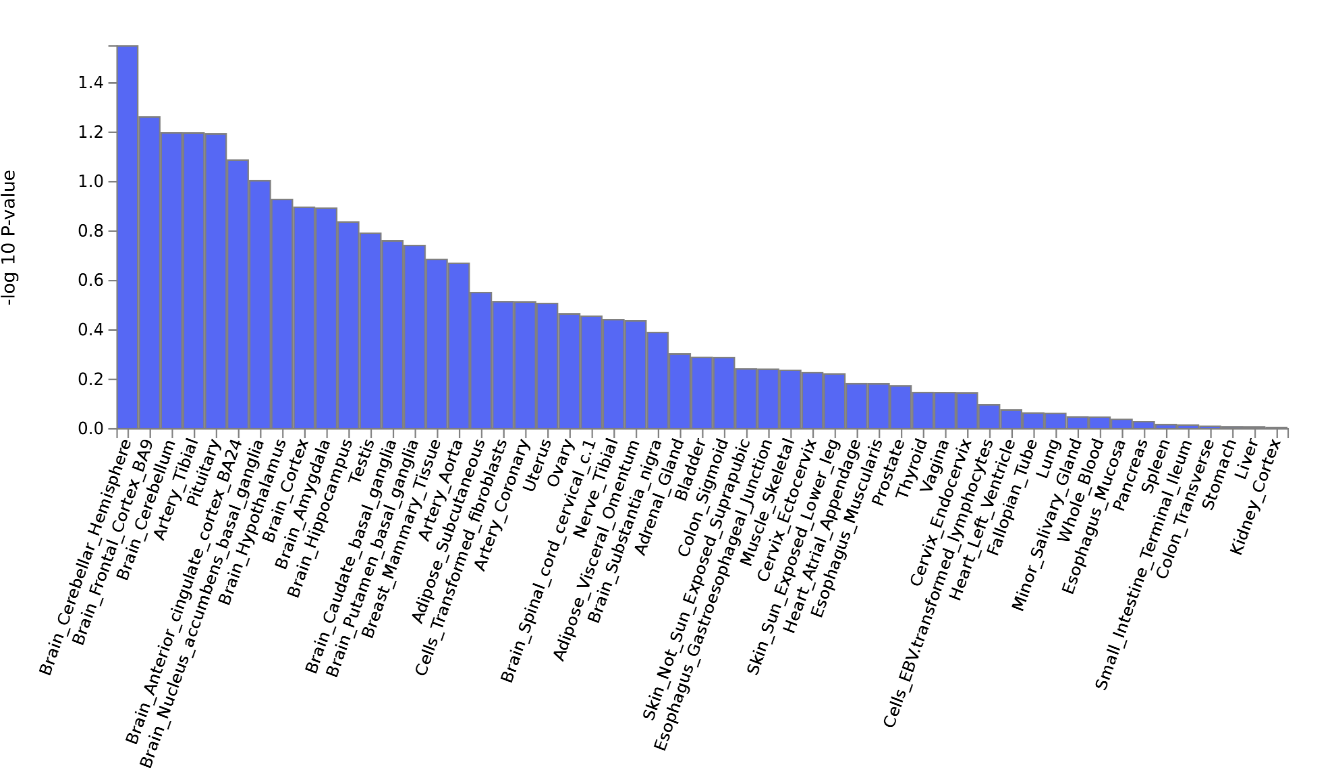

*Tissue specific heritability estimates for SQ-R as provided within FUMA. Tissue expression was constructed using GTEx. General tissue-specific heritability estimates (top), and specific tissue-specific heritability estimates (bottom). Height of the bar represents significance. None of the heritability estimates were significant after Bonferroni correction.*

### SQ-R and its correlates

The SQ-R is self-report measure of systemizing drive, or interest in rule-based patterns^1^, and taps into a variety of domains of systemizing such as interest in mechanical (e.g., car engines), abstract (e.g., mathematics), natural (e.g., the weather), motor (e.g., knitting), and collectible (e.g., stamp collecting) systems.

The idea that systemizing is central to autism was noted in the earliest reports describing autism. In his 1944 paper describing autism^2^, Hans Asperger noted a proclivity for patterns and order in autistic children. Of one child, he writes, “He orders his facts into a system and forms his own theories even if they are occasionally abstruse.” He observes that another child had “specialised technological interests and knew an incredible amount about complex machinery,” while a third child “was preoccupied by numbers.” In Leo Kanner’s 1943 paper, he writes that the children with autism have “precise recollection of complex patterns and sequences^3^.”

These initial clinical observations have been quantified using different measures. For example, on a self-report measure of systemizing (the Systemizing Quotient – Revised, or the SQ-R)^1^, autistic adults, on average, score significantly higher than non-autistic individuals^1,4^. The same pattern of results is seen in autistic children, using the parent-report version of the SQ^5^. More recently, in a dataset of 671,606 individuals including 36,648, we confirmed this observation. Participants completed a short (10-item) version of the SQ-R. We observe, on average, a significant shift towards higher scores on the systemizing in autistic individuals, irrespective of sex (Males, case-control difference: Cohen’s D = 0.3, P < 2.2x10^-16^, females, Cohen’s d = 0.39, P < 2.2x10^-16^). In this large sample, the SQ was significantly and positively correlated with autistic traits measured using the 10-item version of the Autism Spectrum Quotient (AQ) (r = 0.41, P < 2.2x10^-16^), and significantly and positively correlated with sensory difficulties measured using the 10-item version of the Sensory Perception Questionnaire (SPQ) (r = 0.47, P < 2.2x10^-16^). The SQ and the EQ combined explained 41% of the variance in the AQ. In a smaller dataset (811 autistic individuals and 3,906 controls), we identify higher effect size in case-control difference for the full version of the SQ-R that’s used in the GWAS (Males: Cohen’s d = 0.44 , P < 0.01; Females: Cohen’s d = 0.70, P < 0.01)^6^.

Items on the SQ-R measure two of the four DSM-5 criteria in the non-social domain for autism: circumscribed interests ( “Highly restricted, fixated interests that are abnormal in intensity or focus (e.g, strong attachment to or preoccupation with unusual objects, excessively circumscribed or perseverative interest).” ) and sameness/routine (“Insistence on sameness, inflexible adherence to routines, or ritualized patterns or verbal nonverbal behavior (e.g., extreme distress at small changes, difficulties with transitions, rigid thinking patterns, greeting rituals, need to take same route or eat food every day)”).

Examples of items that measure circumscribed interests from the SQ-R include:

1. I can remember large amounts of information about a topic that interests me e.g. flags of the world, airline logos.
2. I have a large collection e.g. of books, CDs, videos etc.
3. When I was young I did not enjoy collecting sets of things e.g. stickers, football cards etc.
4. When I learn about a new category, I like to go into detail to understand the small differences between different members of that category.
5. When I like something, I like to collect a lot of different examples of that type of object, so I can see how they differ from each other.

Examples of items that measure sameness and routine from the SQ-R include:

1. I do not find it distressing if people who live with me upset my routine (negatively coded)
2. When I have a lot of shopping to do, I like to plan which shops I am going to visit and in what order.
3. It does not bother me if things in the house are not in their proper place (negatively coded)
4. I find it difficult to learn my way around a new city.

Several items on the SQ-R map onto items in the Adult Autism Spectrum Quotient (AQ). This is supported by an analysis of the AQ, the EQ and the SQ-R^7^. Combining all three measures, the authors identified a two-factor solution, with the SQ-R and the attention to detail domain in the AQ loading onto a latent non-social factor, and the EQ and the social interaction subscale load onto a latent social factor. The two-factor results remained the same across three groups (N = 363 autistic individuals, N = 439 parents of autistic individuals, and N = 232 controls). Taken together, this suggests that the SQ is an integral part of autistic traits, and is correlated specifically with the non-social domain of the AQ.

**Mapping items on the AQ to items on the SQ-R and items on the EQ**

| Item on the AQ | Equivalent item(s) in the SQ-R | | Equivalent item(s) in the EQ | |
| --- | --- | --- | --- | --- |
| I prefer to do things with others rather than on my own. | X | X | X | X |
| I prefer to do things the same way over and over again. | X | X | X | X |
| If I try to imagine something, I find it very easy to create a picture in my mind. | X | X | X | X |
| I frequently get so strongly absorbed in one thing that I lose sight of other things. | X | X | X | X |
| I often notice small sounds when others do not. | X | X | X | X |
| I usually notice car number plates or similar strings of information. | When I read the newspaper, I am drawn to tables of information, such as football league scores or stock market indices. | I am not interested in the details of exchange rates, interest rates, stocks and shares. | X | X |
| Other people frequently tell me that what I’ve said is impolite, even though I think it is polite. | X | X | I am very blunt, which some people take to be rudeness, even though this is unintentional. | Other people often say that I am insensitive, though I don’t always see why. |
| When I’m reading a story, I can easily imagine what the characters might look like. | X | X | X | X |
| I am fascinated by dates. | When I learn about historical events, I do not focus on exact dates. | X | X | X |
| In a social group, I can easily keep track of several different people’s conversations. | X | X | X | X |
| I find social situations easy. | X | X | I find it hard to know what to do in a social situation. | I don’t consciously work out the rules of social situations. |
| I tend to notice details that others do not. | When I look at a piece of furniture, I do not notice the details of how it was constructed. | When I read something, I always notice whether it is grammatically correct. | X | X |
| I would rather go to a library than a party. | X | X | X | X |
| I find making up stories easy. | X | X | X | X |
| I find myself drawn more strongly to people than to things. | X | X | X | X |
| I tend to have very strong interests which I get upset about if I can’t pursue. | X | X | X | X |
| I enjoy social chit-chat. | X | X | X | X |
| When I talk, it isn’t always easy for others to get a word in edgeways. | X | X | X | I can easily tell if someone else wants to enter a conversation. |
| I am fascinated by numbers. | In maths, I am intrigued by the rules and patterns governing numbers. | X | X | X |
| When I’m reading a story, I find it difficult to work out the characters’ intentions. | X | X | I am good at predicting what someone will do. | I find it easy to put myself in somebody else’s shoes. |
| I don’t particularly enjoy reading fiction. | X | X | X | X |
| I find it hard to make new friends. | X | X | Friendships and relationships are just too difficult, so I tend not to bother with them. | X |
| I notice patterns in things all the time. | When I look at a piece of furniture, I do not notice the details of how it was constructed. | When I listen to a piece of music, I always notice the way it’s structured. | X | X |
| I would rather go to the theatre than a museum. | X | X | X | X |
| It does not upset me if my daily routine is disturbed. | I do not find it distressing if people who live with me upset my routines. | X | X | X |
| I frequently find that I don’t know how to keep a conversation going. |  |  |  |  |
| I find it easy to “read between the lines” when someone is talking to me. | X | X | I can pick up quickly if someone says one thing but means another. | X |
| I usually concentrate more on the whole picture, rather than the small details. | When I listen to a piece of music, I always notice the way it’s structured. | When I look at a painting, I do not usually think about the technique involved in making it. | X | X |
| I am not very good at remembering phone numbers. | X | X | X | X |
| I don’t usually notice small changes in a situation, or a person’s appearance. | X | X | X | X |
| I know how to tell if someone listening to me is getting bored. | X | X | I can easily tell if someone else is interested or bored with what I am saying. | X |
| I find it easy to do more than one thing at once. | X | X | X | X |
| When I talk on the phone, I’m not sure when it’s my turn to speak. | X | X | I can easily tell if someone else wants to enter a conversation. | X |
| I enjoy doing things spontaneously. | X | X | X | X |
| I am often the last to understand the point of a joke. | X | X | X | X |
| I find it easy to work out what someone is thinking or feeling just by looking at their face. | X | X | Other people tell me I am good at understanding how they are feeling and what they are thinking. | I can tune into how someone else feels rapidly and intuitively. |
| If there is an interruption, I can switch back to what I was doing very quickly. | X | X | X | X |
| I am good at social chit-chat. | X | X | X | X |
| People often tell me that I keep going on and on about the same thing. | X | X | X | X |
| When I was young, I used to enjoy playing games involving pretending with other children. | X | X | X | X |
| I like to collect information about categories of things (e.g. types of car, types of bird, types of train, types of plant, etc.). | When I like something I like to collect a lot of different examples of that type of object, so I can see how they differ from each other. | I can remember large amounts of information about a topic that interests me e.g. flags of the world, airline logos. | X | X |
| I find it difficult to imagine what it would be like to be someone else. | X | X | I find it easy to put myself in somebody else's shoes. | X |
| I like to plan any activities I participate in carefully. | When I have a lot of shopping to do, I like to plan which shops I am going to visit and in what order. | X | X | X |
| I enjoy social occasions. | X | X | I don't tend to find social situations confusing | X |
| I find it difficult to work out people’s intentions. |  |  | Other people tell me I am good at understanding how they are feeling and what they are thinking. | I am good at predicting what someone will do. |
| New situations make me anxious. | I avoid situations which I can not control. | X | X | X |
| I enjoy meeting new people. | X | X | X | X |
| I am a good diplomat. | X | X | I can't always see why someons whould have felt offended by a remark. | X |
| I am not very good at remembering people’s date of birth. | I do not tend to remember people's birthdays (in terms of which day and month this falls). | X | X | X |
| I find it very easy to play games with children that involve pretending. | X | X | X | X |

The SQ-R has minimal correlation with measures of intelligence. In a sample from the Cambridge Autism Research Database, we investigated the correlation between the Raven’s Progressive Matrices Test^8^ and the SQ-R in autistic individuals. The correlations were modest and smaller than the correlations observed in the SSC (385 Autistic females: r = -0.09, P > 0.05; 385 autistic males: r = -0.18, P < 0.001). Additionally, two studies have investigated the correlation between the SQ-R and measures of IQ. One study in N = 100 individuals did not find a significant correlation between Baddeley’s Reasoning Task and the SQ-R (r = 0.117, P > 0.05)^9^. In another study, there was no correlation between the SQ-R and full scale IQ in 112 individuals (r = 0.06, P > 0.05)^10^.

### The relationship between social autistic traits and autism

We use four measures of social autistic traits: the EQ, SCDC, the friendship satisfaction, and the family relationship satisfaction scales. Together, these cover several aspects of the social domain of autism as reported by the DSM-5. The DSM-5 criteria have the following three aspects for social difficulties:

1. Deficits in social-emotional reciprocity, ranging, for example, from abnormal social approach and failure of normal back-and-forth conversation; to reduced sharing of interests, emotions, or affect; to failure to initiate or respond to social interactions.

2. Deficits in nonverbal communicative behaviors used for social interaction, ranging, for example, from poorly integrated verbal and nonverbal communication; to abnormalities in eye contact and body language or deficits in understanding and use of gestures; to a total lack of facial expressions and nonverbal communication.

3. Deficits in developing, maintaining, and understanding relationships, ranging, for example, from difficulties adjusting behavior to suit various social contexts; to difficulties in sharing imaginative play or in making friends; to absence of interest in peers.

Two of these (criteria 1 and 3) are captured by these phenotypes, Specifically:

Criteria 1:

From the EQ:

- I can easily tell if someone else wants to enter a conversation.
- I find it hard to know what to do in a social situation.
- I don’t consciously work out the rules of social situations.

From the SCDC:

- Does not seem to understand social skills, e.g. persistently interrupts conversations
- Does not realise if s/he offends people with her/his behavior

Criteria 3:

Items from the EQ:

- It doesn’t bother me too much if I am late meeting a friend.
- Friends usually talk to me about their problems as they say that I am very understanding.
- Friendships and relationships are just too difficult, so I tend not to bother with them.
- I tend to get emotionally involved with a friend’s problems.

From the friendship satisfaction questionnaire:

- In general how satisfied are you with your friendships?

Further, Table 1 in the section above provides the relationship between the EQ and the AQ.

### Standardized SQ-R regression estimates

Here, we outline the method used for standardizing the SQ-R regression estimates, and, as an extension, calculation of the variance explained. We chose to standardize the estimates of SQ-R to make them comparable to the standardized regression estimates for educational attainment. This provides a uniform scale on which GWAS for both the phenotypes were conducted, lending them to analyses using GWIS^11^.

In linear regression, standardized estimates can be obtained from non-standardized estimates using the following formula^12^:

$$\hat{B_{std}}= \frac{\hat{B}\sigma_{x}}{\sigma_{y}}$$

Where, $\hat{B_{std}}$ is the standardized estimate of the regression coefficient, $\hat{B}$ is the non-standardized estimate of the regression coefficient, $\sigma_{x}$ is the standard deviation for the independent variable, and $\sigma_{y}$ is the standard deviation for the independent variable. In the GWAS analyses, y is the phenotype (SQ-R), and x is the genotype. We know $\sigma_{y}$ and $\hat{B}$.

However, $\sigma_{x}= \sqrt{2(MAF)(1-MAF)}$, which has been derived before^13,14^, but we shall derive again below.

$$\sigma_{x}= \sqrt{\sigma_{x}^{2}}$$

$$\sigma_{x}^{2}= \Sigma( x- \bar{x})$$

Let’s assume that the genotype frequencies are in Hardy-Weinberg equilibrium. Let $x_{i}$= 0,1, and 2 for the three genotypes.

P($x_{i}$ = 0) = q^2^

P($x_{i}$ = 1) = 2pq

P($x_{i}$ = 2) = p^2^

$\bar{x}= q^{2}(x_{i}$ = 0) + 2pq($x_{i}$ = 1) + p^2^($x_{i}$ = 2)

= 0 + 2p(1-p) + 2 p^2^

= 2p

$$\sigma_{x}^{2}= q^{2} \left( 0-2p \right)^{2}+2pq \left( 1-2p \right)^{2}+p^{2}\left( 2-2p \right)^{2}$$

$$=2\left( 1-p \right)p$$

$=2(1-MAF)(MAF)$

Therefore,

$$\sigma_{x}= \sqrt{2(MAF)(1-MAF)}$$

And hence,

$$\hat{B_{std}}= \frac{\hat{B}\sigma_{x}}{\sigma_{y}}$$

The variance explained per SNP is R^2^ = ${\hat{B_{std}}}^{2}$

Therefore,

$$R^{2}= \frac{\hat{B}^{2}{\sigma_{x}}^{2}}{{\sigma_{y}}^{2}}$$

### Results of the Genomic SEM analyses

*Results of the genomic SEM analysis with autism, educational attainment, and SQ-R*

V1 = autism, V2 = educational attainment, V3 = SQ-R

| Equation | Unstandardized Estimate | Unstandardized SE | Standardized Est | Standardized SE |
| --- | --- | --- | --- | --- |
| V1 ~ V2 | 0.174344 | 0.040606 | 0.170506 | 0.039712 |
| V1 ~ V3 | 0.240785 | 0.071756 | 0.235523 | 0.070188 |
| V2 ~~ V3 | 0.015051 | 0.003849 | 0.133373 | 0.034106 |
| V1 ~~ V1 | 0.106729 | 0.010321 | 0.904745 | 0.087492 |
| V2 ~~ V2 | 0.112829 | 0.004034 | 1 | 0.035757 |
| V3 ~~ V3 | 0.112866 | 0.011083 | 1 | 0.098197 |

*Results of the genomic SEM analysis with autism, cognitive aptitude, and SQ-R*

V1 = autism, V2 = cognitive aptitude, V3 = SQ

| Equation | Unstandardized Estimate | Unstandardized SE | Standardized Est | Standardized SE |
| --- | --- | --- | --- | --- |
| V1 ~ V2 | 0.123799 | 0.045274 | 0.156563 | 0.057255 |
| V1 ~ V3 | 0.234433 | 0.073881 | 0.22931 | 0.072267 |
| V2 ~~ V3 | 0.026987 | 0.00675 | 0.184938 | 0.046255 |
| V1 ~~ V1 | 0.107305 | 0.010041 | 0.909626 | 0.085114 |
| V2 ~~ V2 | 0.188667 | 0.010237 | 1 | 0.054259 |
| V3 ~~ V3 | 0.112866 | 0.011083 | 1 | 0.098197 |

### Power calculations for polygenic score regression analyses

We conducted power analyses for polygenic score regression, to investigate the statistical power of polygenic scores on the SQ-R to be significantly associated with scores on the RBS-R and ADOS -social and communication domains. Power calculations were conducted using <https://eagenetics.shinyapps.io/power_website/_w_53bbfbeb/_w_70b83796/index.Rmd>, which runs on a previously described theoretical framework^51^. Power calculations depend on multiple parameters including SNP heritability of both the training and the testing phenotypes and genetic covariance between the phenotypes. Genetic covariance can be derived from the genetic correlation and the SNP heritability. In an infinitely large sample, the genetic correlation is equivalent to the square root of the variance explained by the polygenic scores (R^2^). Here, we estimate power calculations at various SNP heritabilities for the second trait and various genetic correlations (graph below). SNP heritability for the SQ-R was 11%. The SNP heritability for the RBS-R was estimated at 15%, and the SNP heritability of ADOS-G was estimated at 26%, based on GCTA GREML calculations.

For the entire sample i.e. SSC + AGRE + LEAP + Paris, we have > 80% power to identify significant effects of polygenic scores on RBS-R at genetic correlations > 0.7. In contrast, for the ADOS social and communication domain, we have > 80% power to identify significant effects of polygenic scores at genetic correlations > 0.55.

### List of authors and their affiliations for the iPSYCH-Broad autism group

| **Group/sample source** | **EMAIL** | **Surname** | **First name** | **Middle name/Initials** | **Professional degrees** | **Affiliations** |
| --- | --- | --- | --- | --- | --- | --- |
| iPSYCH | | Agerbo | Esben |  | DMSci | The Lundbeck Foundation Initiative for Integrative Psychiatric Research, iPSYCH, Denmark ; National Centre for Register-Based Research, Aarhus University, Aarhus, Denmark ; Centre for Integrated Register-based Research, Aarhus University, Aarhus, Denmark |
| iPSYCH | | Als | Thomas | Damm | PhD | The Lundbeck Foundation Initiative for Integrative Psychiatric Research, iPSYCH, Denmark ; Centre for Integrative Sequencing, iSEQ, Aarhus University, Aarhus, Denmark ; Department of Biomedicine - Human Genetics, Aarhus University, Aarhus, Denmark |
| Broad/MHG | | Belliveau | Rich |  |  | Stanley Center for Psychiatric Research, Broad Institute of Harvard and MIT, Cambridge, Massachusetts, USA |
| iPSYCH | | Bybjerg-Grauholm | Jonas |  |  | The Lundbeck Foundation Initiative for Integrative Psychiatric Research, iPSYCH, Denmark ; Center for Neonatal Screening, Department for Congenital Disorders, Statens Serum Institut, Copenhagen, Denmark |
| iPSYCH | | Bækved-Hansen | Marie |  |  | The Lundbeck Foundation Initiative for Integrative Psychiatric Research, iPSYCH, Denmark ; Center for Neonatal Screening, Department for Congenital Disorders, Statens Serum Institut, Copenhagen, Denmark |
| iPSYCH | | Børglum | Anders | D. | MD, PhD | The Lundbeck Foundation Initiative for Integrative Psychiatric Research, iPSYCH, Denmark ; Centre for Integrative Sequencing, iSEQ, Aarhus University, Aarhus, Denmark ; Department of Biomedicine - Human Genetics, Aarhus University, Aarhus, Denmark |
| Broad/MHG | | Cerrato | Felecia |  |  | Stanley Center for Psychiatric Research, Broad Institute of Harvard and MIT, Cambridge, Massachusetts, USA |
| Broad/MHG | | Chambert | Kimberly |  |  | Stanley Center for Psychiatric Research, Broad Institute of Harvard and MIT, Cambridge, Massachusetts, USA |
| Broad/MHG | | Churchhouse | Claire |  | PhD | Analytic and Translational Genetics Unit, Department of Medicine, Massachusetts General Hospital and Harvard Medical School, Boston, USA ; Stanley Center for Psychiatric Research, Broad Institute of Harvard and MIT, Cambridge, Massachusetts, USA ; Program in Medical and Population Genetics, Broad Institute of Harvard and MIT, Cambridge, Massachusetts, USA |
| Broad/MGH | | Daly | Mark | J. | PhD | Analytic and Translational Genetics Unit, Department of Medicine, Massachusetts General Hospital and Harvard Medical School, Boston, USA ; Stanley Center for Psychiatric Research, Broad Institute of Harvard and MIT, Cambridge, Massachusetts, USA ; Program in Medical and Population Genetics, Broad Institute of Harvard and MIT, Cambridge, Massachusetts, USA |
| iPSYCH | | Demontis | Ditte |  | PhD | The Lundbeck Foundation Initiative for Integrative Psychiatric Research, iPSYCH, Denmark ; Centre for Integrative Sequencing, iSEQ, Aarhus University, Aarhus, Denmark ; Department of Biomedicine - Human Genetics, Aarhus University, Aarhus, Denmark |
| Broad/MHG | | Dumont | Ashley |  |  | Stanley Center for Psychiatric Research, Broad Institute of Harvard and MIT, Cambridge, Massachusetts, USA |
| Broad/MHG | | Goldstein | Jacqueline |  |  | Analytic and Translational Genetics Unit, Department of Medicine, Massachusetts General Hospital and Harvard Medical School, Boston, USA ; Stanley Center for Psychiatric Research, Broad Institute of Harvard and MIT, Cambridge, Massachusetts, USA ; Program in Medical and Population Genetics, Broad Institute of Harvard and MIT, Cambridge, Massachusetts, USA |
| iPSYCH | | Grove | Jakob |  | PhD | The Lundbeck Foundation Initiative for Integrative Psychiatric Research, iPSYCH, Denmark ; Centre for Integrative Sequencing, iSEQ, Aarhus University, Aarhus, Denmark ; Department of Biomedicine - Human Genetics, Aarhus University, Aarhus, Denmark ; Bioinformatics Research Centre, Aarhus University, Aarhus, Denmark |
| iPSYCH | | Hansen | Christine | S. |  | The Lundbeck Foundation Initiative for Integrative Psychiatric Research, iPSYCH, Denmark ; Center for Neonatal Screening, Department for Congenital Disorders, Statens Serum Institut, Copenhagen, Denmark ; Institute of Biological Psychiatry, MHC Sct. Hans, Mental Health Services Copenhagen, Denmark |
| iPSYCH | | Hougaard | David | M. | DMSci | The Lundbeck Foundation Initiative for Integrative Psychiatric Research, iPSYCH, Denmark ; Center for Neonatal Screening, Department for Congenital Disorders, Statens Serum Institut, Copenhagen, Denmark |
| Broad/MHG | | Howrigan | Daniel | P. | PhD | Analytic and Translational Genetics Unit, Department of Medicine, Massachusetts General Hospital and Harvard Medical School, Boston, Massachusetts, USA ; Stanley Center for Psychiatric Research, Broad Institute of Harvard and MIT, Cambridge, Massachusetts, USA |
| Broad/MHG | | Huang | Hailiang |  | PhD | Analytic and Translational Genetics Unit, Department of Medicine, Massachusetts General Hospital and Harvard Medical School, Boston, Massachusetts, USA ; Stanley Center for Psychiatric Research, Broad Institute of Harvard and MIT, Cambridge, Massachusetts, USA |
| Broad/MHG | | Maller | Julian |  | PhD | Stanley Center for Psychiatric Research, Broad Institute of Harvard and MIT, Cambridge, Massachusetts, USA ; Genomics plc, Oxford, UK |
| Broad/MHG | | Martin | Alicia | R. | PhD | Analytic and Translational Genetics Unit, Department of Medicine, Massachusetts General Hospital and Harvard Medical School, Boston, USA ; Stanley Center for Psychiatric Research, Broad Institute of Harvard and MIT, Cambridge, Massachusetts, USA ; Program in Medical and Population Genetics, Broad Institute of Harvard and MIT, Cambridge, Massachusetts, USA |
| Broad/MHG | | Martin | Joanna |  | PhD | Stanley Center for Psychiatric Research, Broad Institute of Harvard and MIT, Cambridge, Massachusetts, USA ; Department of Medical Epidemiology and Biostatistics, Karolinska Institutet, Stockholm, Sweden ; MRC Centre for Neuropsychiatric Genetics & Genomics, School of Medicine, Cardiff University, Cardiff, UK |
| iPSYCH | | Mattheisen | Manuel |  | MD | The Lundbeck Foundation Initiative for Integrative Psychiatric Research, iPSYCH, Denmark ; Centre for Integrative Sequencing, iSEQ, Aarhus University, Aarhus, Denmark ; Department of Biomedicine - Human Genetics, Aarhus University, Aarhus, Denmark |
| Broad/MHG | | Moran | Jennifer |  |  | Stanley Center for Psychiatric Research, Broad Institute of Harvard and MIT, Cambridge, Massachusetts, USA |
| iPSYCH | | Mors | Ole |  | MD, PhD | The Lundbeck Foundation Initiative for Integrative Psychiatric Research, iPSYCH, Denmark ; Psychosis Research Unit, Aarhus University Hospital, Risskov, Denmark |
| iPSYCH | | Mortensen | Preben | Bo | DMSci | The Lundbeck Foundation Initiative for Integrative Psychiatric Research, iPSYCH, Denmark ; Centre for Integrative Sequencing, iSEQ, Aarhus University, Aarhus, Denmark ; National Centre for Register-Based Research, Aarhus University, Aarhus, Denmark ; Centre for Integrated Register-based Research, Aarhus University, Aarhus, Denmark |
| Broad/MGH | | Neale | Benjamin | M. | PhD | Analytic and Translational Genetics Unit, Department of Medicine, Massachusetts General Hospital and Harvard Medical School, Boston, USA ; Stanley Center for Psychiatric Research, Broad Institute of Harvard and MIT, Cambridge, Massachusetts, USA ; Program in Medical and Population Genetics, Broad Institute of Harvard and MIT, Cambridge, Massachusetts, USA |
| iPSYCH | | Nordentoft | Merete |  |  | The Lundbeck Foundation Initiative for Integrative Psychiatric Research, iPSYCH, Denmark ; Mental Health Services in the Capital Region of Denmark, Mental Health Center Copenhagen, University of Copenhagen, Copenhagen, Denmark |
| iPSYCH | | Pallsen | Jonatan |  | PhD | The Lundbeck Foundation Initiative for Integrative Psychiatric Research, iPSYCH, Denmark ; Centre for Integrative Sequencing, iSEQ, Aarhus University, Aarhus, Denmark ; Department of Biomedicine - Human Genetics, Aarhus University, Aarhus, Denmark |
| Broad/MHG | | Palmer | Duncan | S. | PhD | Analytic and Translational Genetics Unit, Department of Medicine, Massachusetts General Hospital and Harvard Medical School, Boston, Massachusetts, USA ; Stanley Center for Psychiatric Research, Broad Institute of Harvard and MIT, Cambridge, Massachusetts, USA |
| iPSYCH | | Pedersen | Carsten | Bøcker | DMSci | The Lundbeck Foundation Initiative for Integrative Psychiatric Research, iPSYCH, Denmark ; National Centre for Register-Based Research, Aarhus University, Aarhus, Denmark ; Centre for Integrated Register-based Research, Aarhus University, Aarhus, Denmark |
| iPSYCH | | Pedersen | Marianne | Giørtz |  | The Lundbeck Foundation Initiative for Integrative Psychiatric Research, iPSYCH, Denmark ; National Centre for Register-Based Research, Aarhus University, Aarhus, Denmark ; Centre for Integrated Register-based Research, Aarhus University, Aarhus, Denmark |
| Broad/MHG | | Poterba | Timothy |  |  | Analytic and Translational Genetics Unit, Department of Medicine, Massachusetts General Hospital and Harvard Medical School, Boston, USA ; Stanley Center for Psychiatric Research, Broad Institute of Harvard and MIT, Cambridge, Massachusetts, USA ; Program in Medical and Population Genetics, Broad Institute of Harvard and MIT, Cambridge, Massachusetts, USA |
| Broad/MHG | | Ripke | Stephan |  | MD, PhD | Analytic and Translational Genetics Unit, Department of Medicine, Massachusetts General Hospital and Harvard Medical School, Boston, Massachusetts, USA ; Stanley Center for Psychiatric Research, Broad Institute of Harvard and MIT, Cambridge, Massachusetts, USA ; Program in Medical and Population Genetics, Broad Institute of Harvard and MIT, Cambridge, Massachusetts, USA ; Department of Psychiatry, Charite Universitatsmedizin Berlin Campus Benjamin Franklin, Berlin, Germany |
| Broad/MHG | | Robinson | Elise | B. | PhD | Analytic and Translational Genetics Unit, Department of Medicine, Massachusetts General Hospital and Harvard Medical School, Boston, Massachusetts, USA ; Department of Epidemiology, Harvard Chan School of Public Health, Boston, Massachusetts, USA |
| Broad/MHG | | Satterstrom | F. | Kyle |  | Analytic and Translational Genetics Unit, Department of Medicine, Massachusetts General Hospital and Harvard Medical School, Boston, USA ; Stanley Center for Psychiatric Research, Broad Institute of Harvard and MIT, Cambridge, Massachusetts, USA ; Program in Medical and Population Genetics, Broad Institute of Harvard and MIT, Cambridge, Massachusetts, USA |
| Broad/MHG | | Stevens | Christine |  |  | Stanley Center for Psychiatric Research, Broad Institute of Harvard and MIT, Cambridge, Massachusetts, USA |
| Broad/MHG | | Turley | Patrick |  | PhD | Analytic and Translational Genetics Unit, Department of Medicine, Massachusetts General Hospital and Harvard Medical School, Boston, Massachusetts, USA ; Stanley Center for Psychiatric Research, Broad Institute of Harvard and MIT, Cambridge, Massachusetts, USA |
| Broad/MHG | | Walters | Raymond |  |  | Analytic and Translational Genetics Unit, Department of Medicine, Massachusetts General Hospital and Harvard Medical School, Boston, Massachusetts, USA ; Stanley Center for Psychiatric Research, Broad Institute of Harvard and MIT, Cambridge, Massachusetts, USA |
| iPSYCH | | Werge | Thomas |  | PhD | The Lundbeck Foundation Initiative for Integrative Psychiatric Research, iPSYCH, Denmark ; Institute of Biological Psychiatry, MHC Sct. Hans, Mental Health Services Copenhagen, Denmark ; Department of Clinical Medicine, University of Copenhagen, Copenhagen, Denmark |

### List of authors and their affiliations for the EU-AIMS LEAP group

**KCL:** Dr Antonia San Jose Caceres, Hannah Hayward, Daisy Crawley, Jessica Faulkner, Jessica Sabet, Claire Ellis, Bethany Oakley, Dr. Eva Loth, Prof. Tony Charman, Prof. Declan Murphy

**University of Cambridge:** Dr Rosemary Holt , Jack Waldman, Jessica Upadhyay:

Nicola Gunby, Dr. Meng-Chuan Lai, Gwilym Renouf, Dr. Amber Ruigrok, Emily Taylor, Dr. Hisham Ziauddeen, Dr. Julia Deakin, Professor Simon Baron-Cohen

**UMC Utrecht:** Sara Ambrosino di Bruttopilo, Sarai van Dijk, Yvonne Rijks, Tabitha Koops, Miriam Douma, Alyssia Spaan, Iris Selten, Maarten Steffers, Anna Ver Loren van Themaat

**Central Institute of Mental Health Mannheim**: Dr. Nico Bast, Dr. Sarah Baumeister,

**Radboudumc:** Dr Larry O’Dwyer, Carsten Bours, Annika Rausch

Dr. Daniel von Rhein, Ineke Cornelissen, Yvette de Bruin,

Maartje Graauwmans

**Karolinska Institutet:** Elzbieta Kostrzewa, Elodie Cauvet, Dr. Kristiina Tammimies, Rouslan Sitnikow

**Institut Pasteur:** Dr. Guillaume Dumas , Dr. Yang-Min Kim

### References

1.	Wheelwright, S. J. *et al.* Predicting Autism Spectrum Quotient (AQ) from the Systemizing Quotient-Revised (SQ-R) and Empathy Quotient (EQ). *Brain Res.* **1079,** 47–56 (2006).

2.	Asperger, H. ‘Autistic psychopathy’ in childhood. in *Autism and Asperger syndrome* (ed. Frith, U.) 37–92 (Cambridge University Press, 1944). doi:10.1017/CBO9780511526770.002

3.	Kanner, L. Autistic disturbances of affective contact. *Nerv. Child J. Psychopathol. Psychother. Ment. Hyg. Guid. Child 2* 217–50 (1943).

4.	Baron-Cohen, S., Richler, J., Bisarya, D., Gurunathan, N. & Wheelwright, S. J. The systemizing quotient: an investigation of adults with Asperger syndrome or high-functioning autism, and normal sex differences. *Philos. Trans. R. Soc. Lond. B. Biol. Sci.* **358,** 361–74 (2003).

5.	Auyeung, B. *et al.* Foetal testosterone and the child systemizing quotient. *Eur. J. Endocrinol.* **155,** S123–S130 (2006).

6.	Baron-Cohen, S. *et al.* Attenuation of typical sex differences in 800 adults with autism vs. 3,900 controls. *PLoS One* **9,** e102251 (2014).

7.	Grove, R., Baillie, A., Allison, C., Baron-Cohen, S. & Hoekstra, R. A. Empathizing, systemizing, and autistic traits: Latent structure in individuals with autism, their parents, and general population controls. *J. Abnorm. Psychol.* **122,** 600–609 (2013).

8.	Raven, J. The Raven’s Progressive Matrices: Change and Stability over Culture and Time. *Cogn. Psychol.* **41,** 1–48 (2000).

9.	Ling, J., Burton, T. C., Salt, J. L. & Muncer, S. J. Psychometric analysis of the systemizing quotient (SQ) scale. *Br. J. Psychol.* **100,** 539–552 (2009).

10.	Escovar, E., Rosenberg-Lee, M., Uddin, L. Q. & Menon, V. The Empathizing-Systemizing Theory, Social Abilities, and Mathematical Achievement in Children. *Sci. Rep.* **6,** 23011 (2016).

11.	Nieuwboer, H. A., Pool, R., Dolan, C. V., Boomsma, D. I. & Nivard, M. G. GWIS: Genome-Wide Inferred Statistics for functions of multiple phenotypes. *Am. J. Hum. Genet.* **99,** 917–927 (2016).

12.	Bring, J. How to standardize regression coefficients. *Am. Stat.* **48,** 209 (1994).

13.	Rietveld, C. A. *et al.* Common genetic variants associated with cognitive performance identified using the proxy-phenotype method. *Proc. Natl. Acad. Sci.* **111,** 13790–13794 (2014).

14.	Okbay, A. *et al.* Genome-wide association study identifies 74 loci associated with educational attainment. *Nature* **533,** 539–542 (2016).

15.	Dudbridge, F. Power and Predictive Accuracy of Polygenic Risk Scores. *PLoS Genet.* **9,** e1003348 (2013).

16.	Colvert, E. *et al.* Heritability of autism spectrum disorder in a UK population-based twin sample. *JAMA Psychiatry* **72,** 415–23 (2015).
